# Increased sampling and intra-complex homologies favor vertical over horizontal inheritance of the Dam1 complex

**DOI:** 10.1101/2022.10.04.510763

**Authors:** Laura E. van Rooijen, Eelco C. Tromer, Jolien J. E. van Hooff, Geert J. P. L. Kops, Berend Snel

## Abstract

Kinetochores connect chromosomes to spindle microtubules to ensure their correct segregation during cell division. Kinetochores of human and yeast are largely homologous, their ability to track depolymerizing microtubules however is carried out by the non-homologous complexes Ska1-C and Dam1-C, respectively. We previously reported the unique anti-correlating phylogenetic profiles of Dam1-C and Ska-C found amongst a wide variety of eukaryotes. Based on these profiles and the limited presence of Dam1-C, we speculated that horizontal gene transfer (HGT) could have played a role in the evolutionary history of Dam1-C.

Here, we present expanded analyses of Dam1-c evolution, using additional genome as well as transcriptome sequences and recently acquired 3D structure data. This analysis revealed a wider and more complete presence of Dam1-C in Cryptista, Rhizaria, Ichthyosporea, CRuMs, and Colponemidia. The fungal Dam1-C cryo-EM structure supports earlier hypothesized intra-complex homologies, which enables the reconstruction of rooted and unrooted phylogenies. The rooted tree of concatenated Dam1-C subunits is statistically consistent with the species tree of eukaryotes, suggesting that Dam1-C is ancient, and that the present-day phylogenetic distribution is best explained by multiple, independent losses and no HGT was involved. Furthermore, we investigated the ancient origin of Dam1-C via profile-profile searches. Homology among eight out of the ten Dam1-C subunits suggested that the complex largely evolved from a single multimerizing subunit that diversified into an octameric core via stepwise duplication and sub-functionalization of the subunits before the origin of the Last Common Ancestor.

**Significance:** The Dam1 complex has a crucial role in cell division yet its distribution across species is very patchy. To resolve the evolutionary origin of this peculiar distribution, we used the recently acquired 3D structure to obtain a rooted phylogeny. This study makes an important step in discovering the evolutionary history of the Dam1 complex, by determining that Dam1-C was part of the Last eukaryotic Common Ancestor and arose via stepwise duplications during the transition from prokaryotes to eukaryotes.

## Introduction

During eukaryotic cell division, duplicated sister chromatids are equally divided by the microtubule-based spindle apparatus. Microtubules connect to chromatin via kinetochores, large protein structures that assemble onto centromeric DNA and that regulate equal separation into daughter cells (Cheeseman 2014). The final stages of chromosome segregation requires kinetochores to hold on to depolymerizing microtubule. In yeasts, this role is performed by the Dam1 complex (Dam1-C), which interacts with the Ndc80 complex at kinetochores and forms a ring around the microtubules(Wang et al. 2007). In *Saccharomyces cerevisiae*, Dam1-C is essential and consist of 10 subunits: Dad1-4, Dam1, Duo1, Ask1, Hsk3, Spc34, and Spc19 (Cheeseman, Brew, et al. 2001; Cheeseman, Enquist-Newman, et al. 2001; Enquist-Newman et al. 2001). In *Schizosaccaromyces pombe* Dam1-C has the same function as in *S. cerevisiae*, but is not essential for the short term viability, however, mutations in the subunits do lead to increased chromosome mis-segregations (Thakur & Sanyal 2011).

Dam1-C has an enigmatic evolutionary history. It is the only complex of the yeast kinetochore that has a non-homologous functional counterpart in humans. In humans and other animals the role of Dam1-C is carried out by the non-homologous three-subunit Ska complex (Ska-C) (Cheeseman 2014). Orthologs of the subunits of both protein complexes occur across the eukaryotic tree of life in an largely anti-correlating fashion: 75 surveyed genomes contained one or more orthologs of the subunits of Dam1-C and Ska-C, and only seven genomes were predicted to contain orthologs of both complexes (van Hooff et al. 2017; Cipriano 2013). This striking alternating pattern could be indicative of a last eukaryotic common ancestor (LECA) possessing both Dam1-C and Ska-C, followed by independent, reciprocal loss. Alternatively, the pattern could have arisen through horizontal gene transfer (HGT) and displacement of one or both complex(es) across the eukaryotic tree of life. Phylogenetic analyses failed to unequivocally favor either scenario, as crucial branches in the phylogenies of the paralogs had low supports and the phylogeny from the concatenated alignment of the full complex could not be rooted. HGT was ultimately favored over a LECA scenario, as HGT is “simpler”: the LECA scenario may imply both complexes to have had a different function in LECA in order to have co-existed, and to subsequently converge towards a kinetochore function, and, even more improbable, to do so in an alternating and independent manner in different lineages. In addition, it implies more losses and implies that many ancestral branches had both complexes, which contrasts the current-day underrepresentation of species with both.

A recent Cryo-EM structure of *Chaetomium thermophilum* Dam1-C provided new insights (Jenni & Harrison 2018): This structure revealed that Dam1-C consists of two arms each formed by five-helix bundles, with the N-termini of the subunits at the distal ends of the arms (Figure 1). From this structure, the authors perceived each subunit has one “structural paralog” in the other arm, leading to five pairs: Ask1-Dad3, Dad2-Duo1, Dad4-Dad1, Hsk3-Dam1 and Spc19-Spc34. These putative paralogous relationships confirmed two earlier deep homologies inferred from profile-vs-profile comparisons (van Hooff et al. 2017) and the suggested three additional homologous pairs. Besides the implied large role for duplication in the history of the Dam1-C, the symmetrical structure of the Dam1-C in principle allows for a concatenation of the subunits of each separate arm and thereby infer a rooted tree – the root can be placed between the arms. Such a rooted phylogeny would enable to discern patterns of inheritance, including HGT, in contrast to the previously published, unrooted phylogeny based on an all-subunit concatenation.

**Fig. 1:**
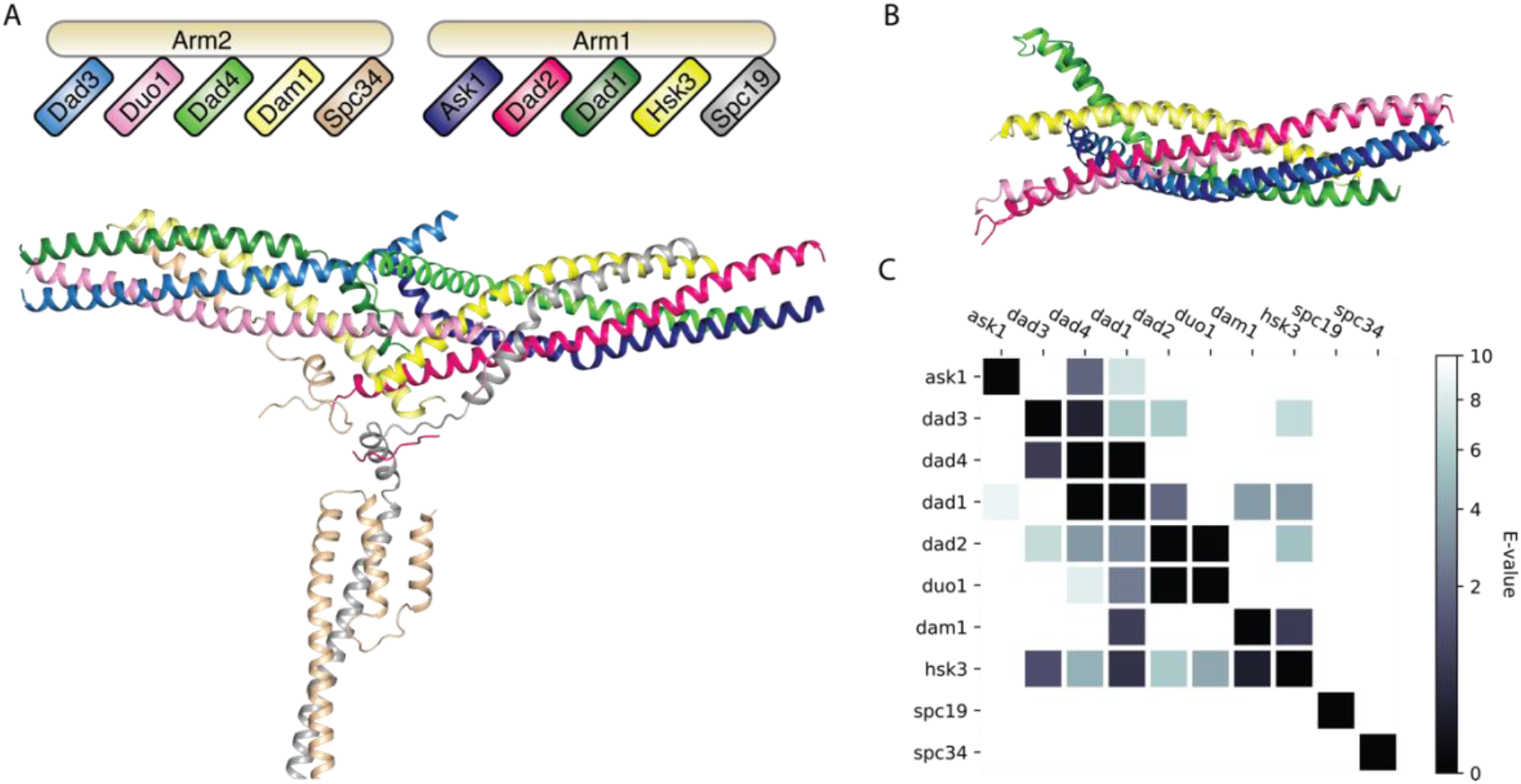
Dam1-C contains four paralogous pairs. A) structure of Dam1-C, the subunits are color-coded according to the legend, B) the structural alignment of Dam1-C subunits to their paralogs, and the subunits are color-coded as in the legend of A, C) heatmap of the profile vs profile hits for the separate subunits of the Dam1-C (cutoff is e-value=10).

Next to the 3D structure, new transcriptome and genome data are becoming available rapidly and specifically relevant for DAM1-C are under-sampled groups such as CruMs, Cryptista and Colponemidia. In addition to further uncovering the phylogenetic distribution of both complexes, these new data could aid in resolving their evolutionary histories. We utilize this novel sequence and structural information to improve our understanding of the evolutionary history of Dam1-C, including reconstructing a rooted phylogeny for Dam1-C. Our analyses suggest Dam1-C was already part of the LECA and indicate that this complex evolved through multiple intra-complex duplications.

## Results

### New transcriptomes reveal near-complete Dam1-C presences in multiple lineages outside fungi

To revisit the evolutionary history of Dam1-C, a set of 181 eukaryotic species was compiled. This set expanded a previous data set in order to make use of predicted proteomes at phylogenetic positions that are relevant for Dam1-C and Ska-C (Deutekom et al. 2021) (Figure 2, supplementary figure 1). The most important additions were as followed: First, four Ichthyosporea were added to better investigate the presence of Dam1-C in deep branching Holozoa (Grau-Bové et al. 2017). Second, early-branching species within clades without known Dam1-C presence were added, including *Mantamonas plastica* and *Rigifila ramosa* (CRuMs), two colponemidia (Alveolata) and *Andalucia godoyi* (Jakobida). Third, the data from the Marine Microbial Eukaryotic transcriptome sequence project (MMETSP, version 3(Keeling et al. 2014; Johnson et al. 2019)) was explored to find more species having a near-complete presence of Dam1-C. In the MMETSP, three additional Rhizaria species were found with Dam1-C and for *Bigelowiella natans*, eight subunits instead of the previously detected five subunits were found in the strain designated CCMP1259. Fourth, three new Cryptista species were added to give strength to this group in addition to *Guillardia theta* from previous analyses, as *G. theta* is of interest because it has both Dam1-C and Ska-C. Fourth, an extensive search was performed to add more Rhodophyta with Dam1-C to the dataset, yet only one additional species with Dam1-C subunits was found, namely *Cyanidium caldarium*. Finally, the genome of the filamentous fungus *Chaetomium thermophilum* was added, since this is the organism whose Dam1-C structure was elucidated (Jenni & Harrison 2018)

**Fig. 2:**
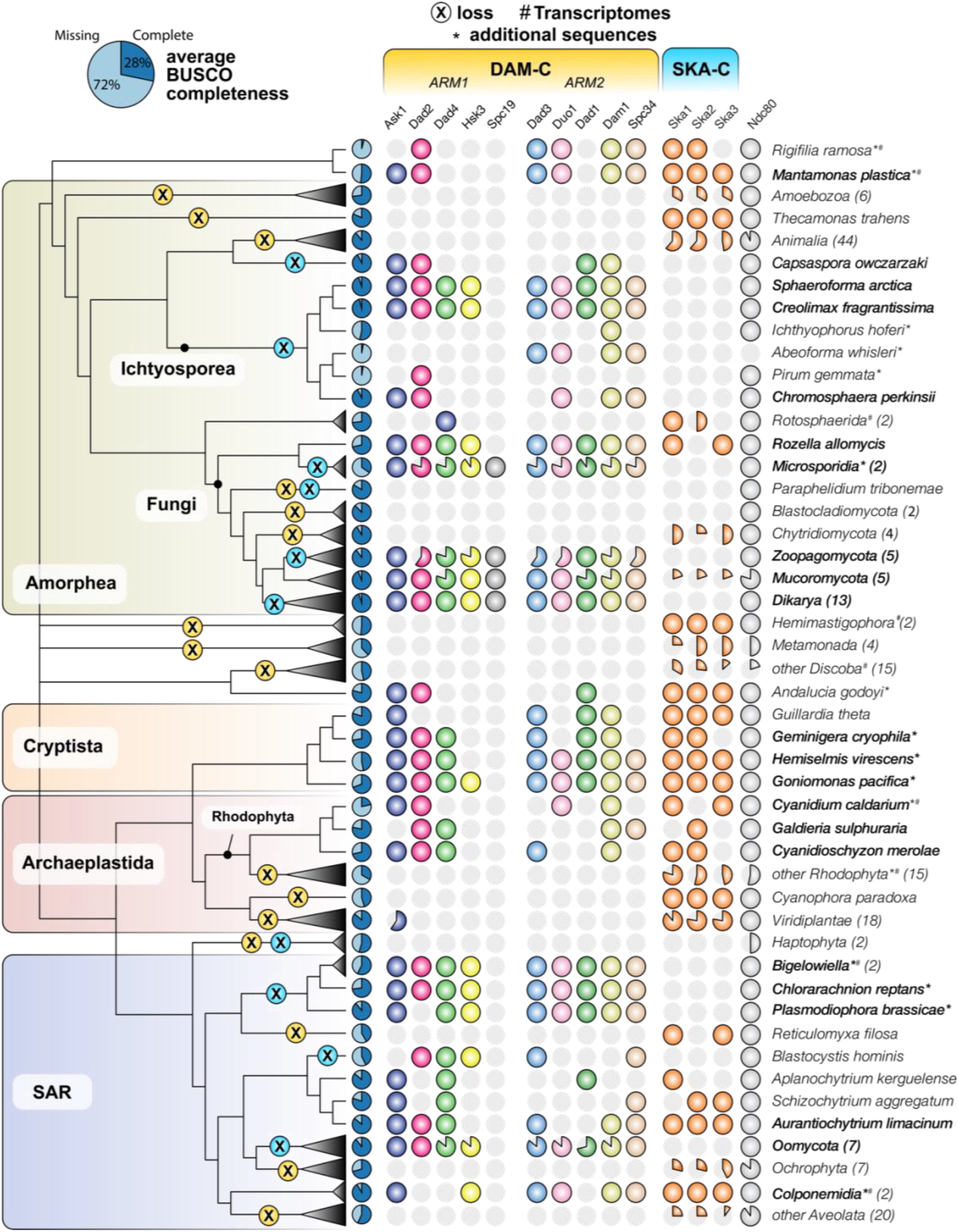
Presence-absence profiles of Dam1-C subunits, Ska-C subunits and Ndc80 across selected eukaryotes. Filled circles denote presences. An asterisk indicates the presence of this species(strain) is a new addition compared to earlier work and a hashtag indicates a transcriptome. The number behind each taxon describes the number of species in our genome set of that taxon. When the pie is not full the section of the pie that is filled shows how many of the members in that group contain that subunit. The eukaryotic groups are color-coded according to the legend. Species in bold indicates the species is part of the tree in figure 3.

Orthologs of ten Dam1-C subunits were detected using Hidden Markov Model (hmm) profile searches against the compiled eukaryotic dataset (see Methods). As expected, a complete Dam1-C complex was found among Fungi (Figure 2). Dam1-C subunits remain undetected in Metazoa, but clear hits were observed in their unicellular relatives, namely the Ichtyosporea (e.g. *Hemiselmis virescens, Abeoforma whisleri, Pirum gemmata, Ichthyophonus hoferi, Chromosphaera perkinsii, Creolimax fragrantissima*, and *Spaeroforma arctica*) and the filasterean *Capsaspora owkzarzaki*. Orthologs of Dam1-C subunits were also detected in *Mantamonas*, which is currently thought to be positioned basal to Amorphea (Fungi, Holozoa, Amoebozoa, and other unicellular relatives) (Burki et al. 2020). Outside the Amorphea, a near-complete Dam1-C was found in Cryptista, Rhizaria, Stramenopila, and Jakobida. Five orthologous subunits of the Dam1-C were found in an early-diverging Alveolata branch (Colponemidia). Even though we found near-complete Dam1-C presences outside of Fungi, one Dam1-C subunit remained conspicuously absent in non-fungal lineages, namely Spc19, and we therefore deem it Zygomycete and Dikarya-specific.

Comparing the phylogenetic profile of Ska-C (Figure 2) to that of Dam1-C using this expanded species set provides the following picture. It confirmed previous observations of a complementary phylogenetic distribution of Dam1-C and Ska-C. Most Ndc80-containing species (76%) have either Dam1-C (64/165) or Ska-C (92/165), and only 8% (14/165) of the species have both complexes. Dam1-C subunits were found in six out of nine eukaryotic major groups included in this study (CRuMs, Amorphea, Discoba, Metamonada, TSAR, Cryptista, and Archaeplastida) and Ska-C in eight out of nine (CRuMs, Amorphea, Discoba, Metamonada, Hemimastigophora, TSAR, Cryptista, and Archaeplastida). Thus, although the occurrence of Dam1-C is more widespread than previously estimated (e.g., present in Mantamonas and early alveolates) and more complete in two lineages where it was observed before (from three to nine subunits in Ichthyosporea and from five to nine subunits in Cryptista), the tendency of Ska-C and Dam1-C to be mutually exclusive remains. However, Ska-C’s and Dam1-C’s combined presence in early-branching Alveolata and in an early-branching relative of Fungi, Metazoa and Amoebozoa (CRuMs) make a stronger case for LECA carrying both Ska-C and Dam1-C. In addition, more sequences allowed for a renewed, richer phylogenetic investigation into their evolutionary histories.

### Four paralogous pairs in the Dam1-C

Jenni and colleagues (Jenni & Harrison 2018) suggested that Dam1-C consists of five “structural paralogs” (Figure 1A). Based on Hidden Markov profile-profile comparisons, two sets of paralogous subunits were previously predicted in Dam1-C, namely Duo1-Dad2 and Dad1-Dad4-Ask1. Two of these paralogous pairs were confirmed by the structural paralogs: Duo1-Dad2 and Dad1-Dad4. To validate the proposed putative homologies, the two arms were structurally aligned (Figure 1B). The structures of the subunits Ask1-Dad3, Dad2-Duo1, Dad4-Dad1, and Hsk3-Dam1 can be super-positioned perfectly, as indicated by the low root mean square deviations (RMSD, Ask1-Dad3: 4.900 Å, Dad2-Duo1: 4.714Å, Dad4-Dad1: 1.528 Å, and Hsk3-Dam1: 2.793 Å). Profile-profile searches between the alignments of orthologs of each Dam1-C subunit find as reciprocal best matches three of the four paralogous pairs (Figure 1C). Ask1-Dad3 is the exception, as these subunits are not directly retrieved in the profile-profile search, but they are linked via homology with other subunits (i.e. Dad3-Dad4 and Dad4-Ask1) and can be almost perfectly super-positioned (Figure 1B). The profile searches do not recover any homology between Spc19 and Spc34, which agrees with our failure to structurally align these subunits convincingly (RMSD: 27.318 Å). Here our structural alignments combined with profile based sequence searches confirm that Dam1-C out of 4 homologs pairs.

### The phylogenetic tree of concatenated subunits of both Dam1-C arms is consistent with the species tree

Incongruence of a gene tree with the species tree is the gold standard to distinguish horizontal gene transfer (HGT) from vertical descent. However, individual gene trees of Dam1-C subunits have a poor resolution due to limited phylogenetic signal (Supplementary figure e). In contrast, the concatenated subunit tree of Dam1-C is highly consistent with the eukaryotic phylogeny, indicating vertical descent (Figure 3A, Supplementary figure3). Nevertheless, depending on the root of this tree, it could also indicate an early HGT for example from fungi to other eukaryotic species. The symmetrical paralogy of the structure of Dam1-C allows for concatenation of the subunits from the two separate arms and thereby root the tree between the arms, to study vertical and horizontal signals in the evolution of Dam1-C. For a species to be suited for the rooted phylogenetic tree, we required it to have at least two subunits of the Dam1-C in both arms (Figure 2, species that are in bold). Spc19 and Spc34 were excluded from the concatenated alignment because structural alignments as well as profile-profile searches did not unequivocally reveal homology between these two subunits and because Spc19 is lineage-specific in our homology searches.

**Fig. 3:**
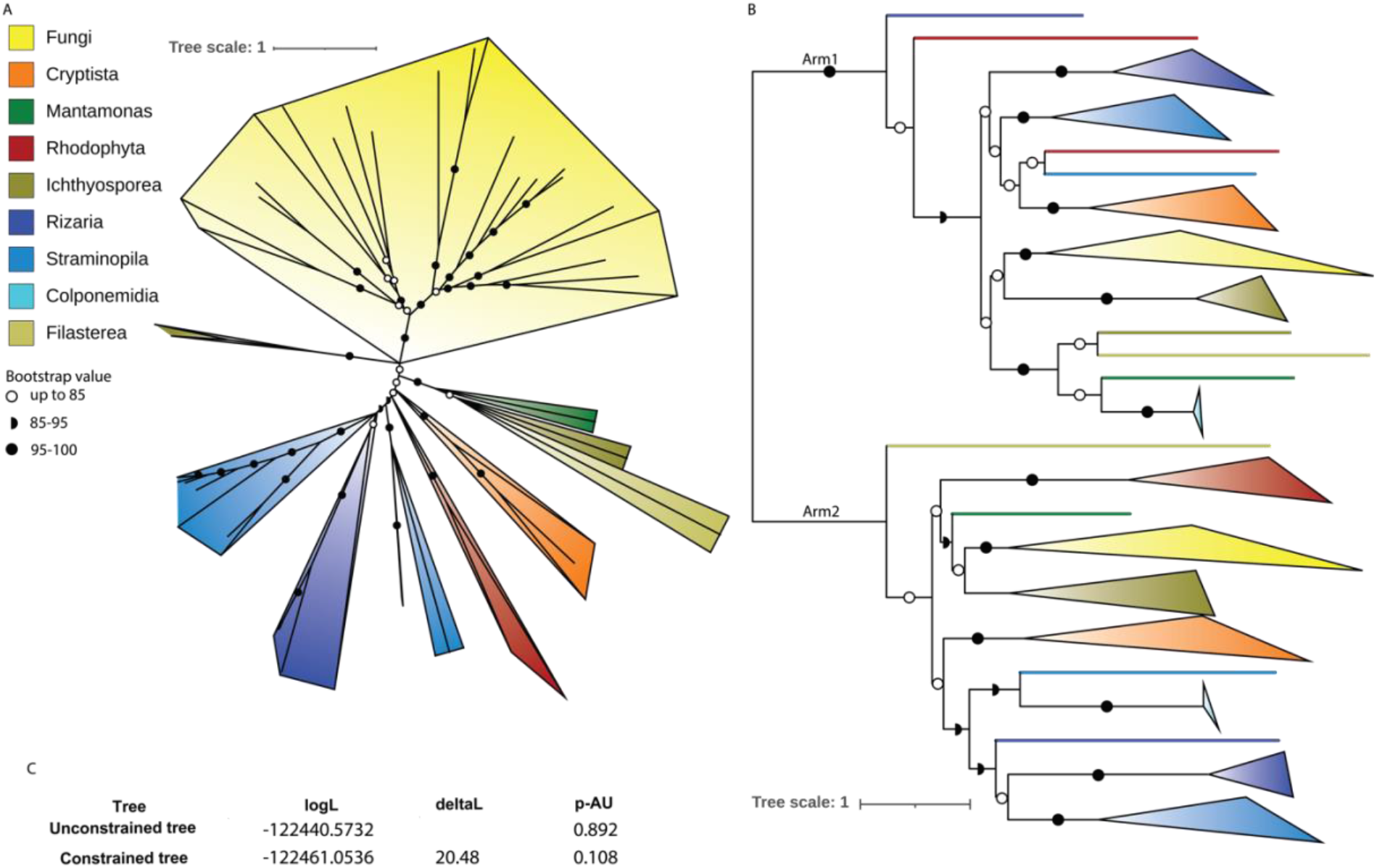
The phylogenetic tree of Dam1-C is consistent with the species tree. A) Phylogeny inferred from concatenating the Dam1-C subunits, the eukaryotic groups are color-coded as in the legend. B) Phylogeny inferred from aligning the subunits pairs and concatenation of the subunits belonging to each arm. C) Topology testing of the rooted concatenated tree and the constrained tree to follow the species tree. LogL is the likelihood of the specific tree. DeltaL is the difference between both trees and p-AU is the topology test. A tree is rejected if p<0.05

When concatenating the subunits per arm, the division between the arms is well supported (Figure 3B, Supplementary figure 4) and retrieves most of the major eukaryotic phyla/taxa as monophyletic clusters, albeit less consistent than the full concatenation. The rooting by the arm-separating internal duplications allows us to see if any eukaryotic taxon could have acted as a donor to all other lineages, which was previously hypothesized for fungi (van Hooff et al. 2017). In this specific hypothesis, one expects fungi to be located at the base of the phylogenetic tree. In both the arm1 and arm2 cluster, fungi are not at the base, making it unlikely that Dam1-C originated in and was transferred from Fungi. The most prominent incongruence with the species tree is *Plasmodiophora brassicae*’s failure to cluster with other Rhizaria. Inspection of the alignment revealed that the likely reason for this is incomplete data (gaps in the genes that are predicted) in *P. brassicae* and other Rhizaria.

Although both arms of the rooted tree are largely consistent with the species tree, they are not identical to the current consensus on the eukaryotic tree of life (Burki et al. 2020). To determine if the data is statistically significantly inconsistent with the consensus species tree, topology testing was used (Approximately Unbiased Test, implemented by IQtree (Nguyen et al. 2015; Shimodaira 2002)) was used. Specifically, we tested whether the tree obtained in figure 3B is significantly different from the species tree by constraining our phylogenetic analysis to obey to the species tree (Supplementary Figure 5), and subsequently comparing the likelihoods of the tree in figure 3B and the constrained tree. The constrained trees are not significantly worse than the unconstrained tree (Figure 3C), revealing that the topology in line with the species tree is an equally valid hypothesis for the evolution of Dam1-C. This supports vertical inheritance as a strong component of Dam1-C evolution.

### Homologies within Dam1-C arms suggest a large role for intra-complex duplications in the ancient origin of Dam1-C

A pre-LECA origin of Dam1-C leaves open the question of how the Dam1-C evolved during the transition from prokaryotes to eukaryotes. Given that profile searches of individual Dam1-C subunits hit other subunits than their closest homolog (Figure 1C), albeit at grey zone e-values, some of the established paralogous pairs could be homologous to one another as well. A profile-versus-profile search using merged alignments of the paralogous pairs indeed indicates homology amongst the paralogous pairs (Figure 4A). The merged alignments of the paralogous pairs generally first hit the separate profiles of the separate subunits, then the profiles of the other paralogous pairs, and finally proteins outside of the complex. The putative deep homology between Dam1, Ask1, Dad1-4, Duo1 and Hsk3 allowed building a phylogeny containing these eight Dam1-C subunits (Figure 4B). The clustering in this phylogeny confirms the closest paralogy relation of Dam1-Hsk3, Dad1-Dad4 and Ask1-Dad3, however further inferences are hampered by the low support values of the internal branches.

**Fig. 4:**
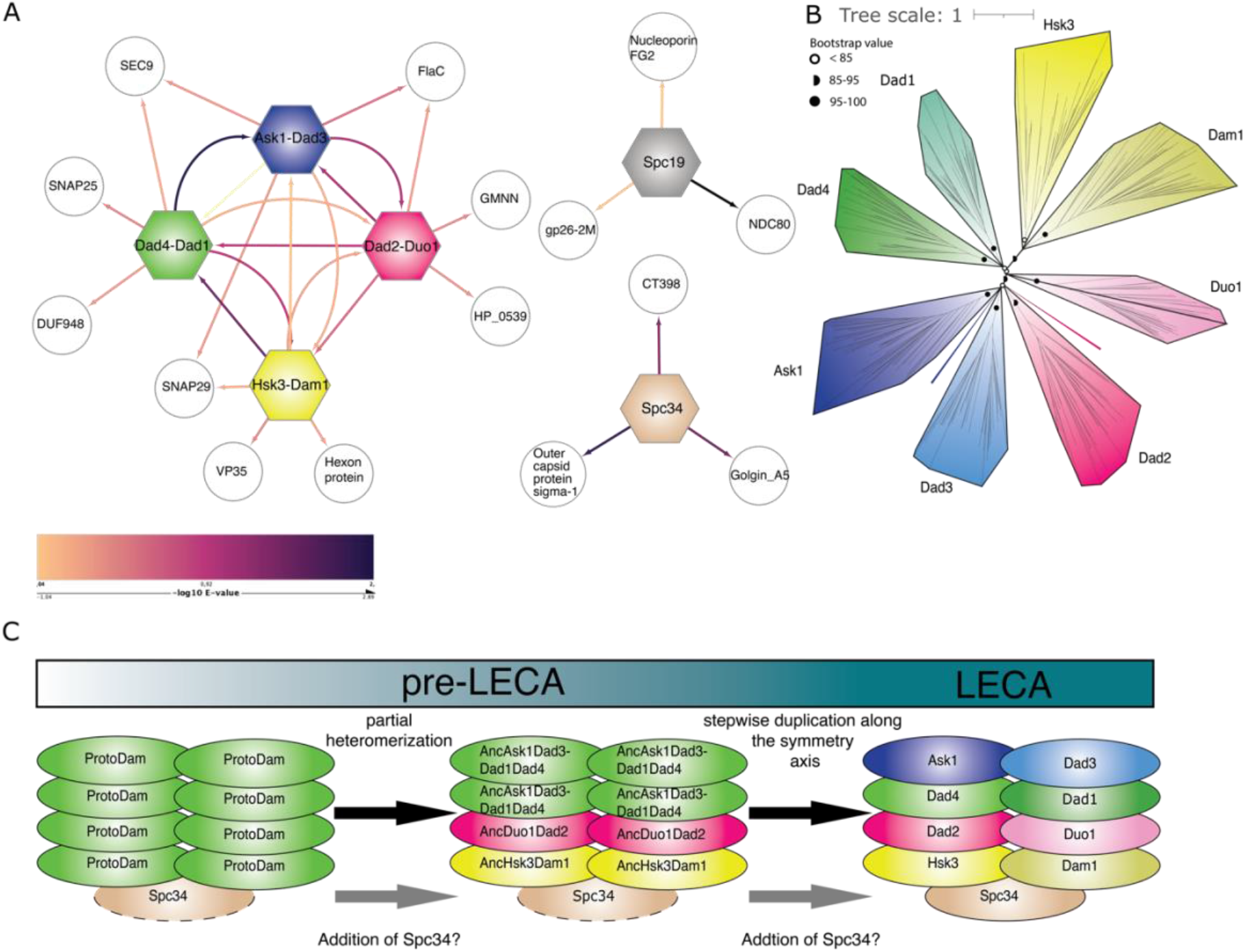
Relation amongst the subunits and a scenario for the origin of Dam1-C. A) The results of the profile vs profile searches are represented as a network. Hexanal nodes represent the profiles of the merged paralogous subunits and the round nodes represent PDB and Pfam profiles. The lines indicate e-values, darker line depict lower e-values. Eight Dam1-C subunits emerge as a consistent cluster B) Phylogeny if the 8 paralogous Dam1-C subunits. Constructed using mafft merge and iqtree, see methods C) Scenario for the origin of the DAM1-C during the transition from prokaryotes to eukaryotes culminating in a 9 subunit complex in LECA.

The homologies within the Dam1-C suggest a large role for intra-complex duplications in the origin of the Dam1-C. We suggest the following scenario based on the protein complex structure, the intra-complex homologies and the phylogeny of subunits (Figure 4C). First, there was a single proto-Dam subunit that homo-multimerized into a Dam1-C like structure. Subsequent (stepwise) duplications resulted in the formation of a heteromer along the parallel axis (Figure 4C, step 2). The next wave of (stepwise) duplications among the bifold symmetry axis and sub-functionalization of the separate subunits, together resulted in an eight-subunit Dam1-C in early eukaryotic evolution. At some point before LECA, Spc34 was added. It could have been recruited when Dam1-C was still a homomer, or much later. Spc19 is most likely a (post-LECA) addition to the complex at the common ancestor of Dikarya and Zygomycetes.

## Discussion

Adding novel lineages (Colponemidia, Mantamonas) and increasing the resolution of previously analyzed lineages (Cryptophyta, Cercozoa) strengthens the unique phylogenetic distribution of Dam1-C as sparse yet wide, and largely anti-correlating with Ska-C. New structural data of Dam1-C (Jenni & Harrison 2018) confirmed previously postulated intra-complex homologies (van Hooff et al. 2017) and allowed inference of an arm-based phylogenies. These phylogenies strongly suggest that Dam1-C evolved through vertical descent, and that the complex was likely present in the last eukaryotic common ancestor (LECA). Such an ancient origin for Dam1-C would mean that the sparse phylogenetic distribution is the result of widespread independent loss, having occurred in at least 14 lineages. This ancient origin also raises the question of the origin of the complex. Deep homologies inferred from merged profile-vs-profile searches and from the 3D structure suggest a scenario where Dam1-C arose through multiple rounds of intra complex duplications, during eukaryogenesis.

An origin by intra complex duplications during eukaryogenesis is has been reconstructed for other complexes, providing precedent for such an inference in the case of Dam1-C. For example, the SM/LSM rings which form a heteroheptameric ring and accompanies snRNA in the spliceosome. The rings consist of seven proteins each, which are all homologous and arose in two distinct waves of duplications. Between these waves of duplications, these copies underwent extensive sequence divergence, which makes determining the precise order of duplications difficult (Veretnik et al. 2009; Scofield & Lynch 2008).

Despite progress on elucidating the deep evolutionary history of Dam1-C, uncertainties remain. For example, it is not clear why the genomes of many organisms seemingly contain only a subset of Dam1-C subunits. The inference of incomplete complexes could be due to data problems (see below) or to the difficulty to find homologies. Alternatively, Dam1-C subunits might be functional even if not all subunits are present, as large-scale studies of protein complexes have suggested (Fokkens & Snel 2009; Schultz & Seidl 2009) or the complex is functional by replacement of missing subunits(Morett et al. 2003). The uncertainty brought about by the partial presence of Dam1-C subunits is especially relevant in the Rhodophyta. Rhodophyta is one of the main lineages with a primary plastid which has spread by eukaryote-to-eukaryote endosymbiosis(Strassert et al. 2021). Although exhaustive searches were performed for Rhodophyta, the absence of any red alga with full Dam1-C prevents us to assess if part of the sparse distribution of Dam1-C can be explained by secondary endosymbiosis (re)introducing Dam1-C into lineages such as Stramenopila or Cryptophyta.

Even though the arm-based tree is statistically consistent with the species tree (as described by Burki(Burki et al. 2020)), more species and more sequences per species could solidify this result. Specifically, the paraphyly of Rhizara caused by erroneous placement of *P. brassicae* is likely to be resolved when Dam1-C subunits from closely related species would be available. Another potential data issue is gene prediction problems. We relied on predicted proteomes and especially missing genes as well as wrongly predicted genes could have negatively impact phylogenetic resolution. The genes encoding Dam1-C subunits are short (average length:102.8 amino acids) and therefore can be more easily missed by gene prediction software (Deutekom et al. 2019). We circumvented this issue by only including species with a minimum of two subunits per arm into the phylogenetic analysis.

Dam1-C joining Ska-C as present in early eukaryotic evolution eliminates the mystery of transferring an entire complex amongst eukaryotes. Instead, it raises another issue; why would LECA have two systems that have the same function and why were Dam1-C and Ska-C lost so often? Some intuition can come from other proteins with anti-correlating patterns as these are not fully unique to Dam1-C and Ska-C. Two examples that also display such patterns and thus could help us to understand Dam1-C and Ska-C are (1) the paralogs elongation factor-I alpha (EF-Iα) and elongation factor-like (EFL)(Cocquyt et al. 2009; Kamikawa et al. 2013) and (2) the paralogs single subunit of ATP citrate lyase (ssACL) and double subunit ACL(Gawryluk et al. 2015). Both pairs were inferred to be present in LECA and their anti-correlating pattern has been attributed to a combination of differential loss and specialization. The differential loss was speculated to stem from the overlapping function of the paralogs, which led one of the two proteins to become progressively less expressed, and subsequently lost in most existing lineages. In addition, especially in the case of EFL, the few species that do contain both proteins, one paralog is retained for a subset of the original functions and this sub-functionalization prevents losing this protein. These two examples are not fully comparable to Dam1-C and Ska1-C since they pertain to paralogs instead of analogs and encompass single proteins instead of protein complexes, nevertheless they provide support for functional redundancy at LECA followed by reciprocal loss and some degree of functional specialization to also be at play for Dam1-C and Ska1-C.

Another explanation is that one of the complexes had a different function. It was recently shown that Dam1-C, in addition to its function in the kinetochore, also has a function in hyphal tip growth in fungi (Shah et al. 2019). This second function invites the hypothesis that in LECA, Ska-C had the kinetochore as the main function and Dam1-C had an additional function in another molecular process where microtubule tracking plays a role, like for example hyphal tip growth in some present-day fungi. Subsequently, loss of the need for this ancestral Dam1-C function in lineages such as Metazoa and Viridiplantae would incite loss of Dam1-C, while a strong need for this ancestral Dam1-C function would allow loss of Ska1-C, provided Dam1-C gained kinetochore activity. This proposed scenario implies that it is not deleterious to possess both Dam1-C and Ska-C (similar to EFL or ALC) but instead that a selection for gene loss from a feature-rich LECA explains the pattern. An additional two-function hypothesis is that the complexes were mitosis- and meiosis-specific, such as is known for Rec8 and Rad21 (Parisi et al. 1999). In current species with both complexes this would predict that one complex (either Dam1-C or Ska-C) has a function in mitosis and the other in meiosis.

Our phylogenomic findings propose that LECA contained both Dam1-C and Ska-C and Dam1-C arose through intra complex duplications. This hypothesis also raises questions as outlined above, which highlight the need for experimental cell biology to study the functional overlap and differentiation in the - mostly understudied - organisms that contain both complexes such as *Rhizophagus irregularis, Guillardia theta* or *Aurantiochytrium limacinum*.

## Methods

### Compiling the proteome database

For studying the presences and absences of subunits of Dam1-C, Ska-C and Ndc80 across the eukaryotic tree of life, a dataset was compiled containing the predicted proteomes, genomes and transcriptomes from 181 eukaryotic organisms from different supergroups: 86 Opisthokonta, 6 Amoebozoa, 26 Archaeplastida, 4 Cryptista, 13 Excavata, 2 Haptophyta, 2 Hemimastigophora, 44 SAR and 1 Apusozoa (Supplementary table 1). The initial set was constructed as described previously (Deutekom et al. 2021).This initial set was expanded using the following criteria: 1. species were selected to represent eukaryotic diversity and allow for a detailed analysis of the evolution of Dam1-C. 2. If available, two species were selected per clade and commonly used model organisms were preferred over other species. 3. If multiple proteomes, genomes or transcriptomes were available for a single species, the one with the most complete Dam1-C complex was selected.

The Cryptista and the extra Rhizaria were obtained from the MMETSP (Marine microbial Eukaryotic transcriptome sequence project version 3(Keeling et al. 2014; Johnson et al. 2019)). To be able to make use of the transcriptomes in MMETSP we translated them using transeq (EMBOSS:6.6.0.0). We replaced *Bigelowiella natans* strain CCMP2755 with strain CCMP1259 compared to van Hooff et al, because for this strain we were able to find 8 subunits instead of 5 subunits. For the Colponemidia, Mantamonas, Ichthyosporea and Rhodophyta, the EukProt database version 2 was used (Richter et al. 2020).

### Orthologs detection

Orthologs of Dam1-C, Ska-C and Ndc80 (fig1) were obtained by use of HMM profiles as constructed by van Hooff et al. 2017 (van Hooff et al. 2017). Although hmmer searches are primarily homology searches, we have previously demonstrated that hmmer models capture a single orthologous subunit per species for Dam1-C subunits. The ‘hmmsearch’ tool from the HMMER package (http://hmmer.org/, HMMER 3.1b1) and the initial profiles from van Hooff et al. were used to search through the updated database to find the subunits in the newly-added species. The profiles were updated by adding the subunits from the new species to the multiple sequence alignment using MAFFT E-INS-i (MAFFT v7.271) and then were created using ‘hmmbuild’ tool from the HMMER package. When searching with these updated profiles, we did not detect any additional Dam1-C subunit orthologs. Due to the structure of the Dam1-C subunits, i.e. coiled-coil structure, the HMMER output was assessed manually. If multiple hits per species were found, phylogenetic analysis was used to differentiate orthologs from paralogs.

From the MMETSP dataset, species were added if hits within the output of this species had an E-value < 10**-3 and if it had 4 or more subunits after manual curation.

### Profile-vs-profile searches

To investigate if all the subunits are homologous to one another, a profile-versus-profile search was preformed (Figure 4A), using the database of Pfam 31, pdb70, a database of profiles of kinetochore proteins and of the merged alignments of the subunit pairs of the Dam1-C. The merged alignments were aligned using MAFFT merge E-INS-i (MAFFT v7.271) and filtered using trimAl gt 0.05. The alignments were manually curated for gene prediction problems. HHsuite (version 3.3.0, tool hhsearch) was used for profile-versus-profile search using the merged protein alignments of the coupled subunits and the separate protein alignments for each subunit. The fasta files used to create the merged profiles can be found in de supplementary files.

### Phylogenetic analysis

For the subunit tree (Figure 4B), the paralogy of the Dam1-C subunits was confirmed by the profile-versus-profile results. The sequences of each orthologous group were aligned using MAFFT einsi (Katoh et al. 2005) (MAFFT v7.271). For building the tree we then aligned the MSAs of the subunits of the Dam1-C by using MAFFT merge E-INS-i. The alignments are trimmed using trimAl (Capella-Gutiérrez et al. 2009) (trimAl v1.4.rev15) with gt 0.05. IQ-TREE was used to perform a model finder and infer the phylogeny as advised by Kalyaanamoorthy et al. (2017)(Kalyaanamoorthy et al. 2017). IQtree was run with a 1000 ultrafast bootstraps and the model selected by ModelFinder was VT-F-R7. The sequences of the orthologous groups can be found in supplementary files, as well as the alignments and the IQ-TREE tree file.

### Concatenated tree

For the concatenated tree based on the arms in the structure of Dam1-C, the subunits were aligned to their structural paralog (Figure 3B). These alignments were concatenated to create a phylogeny. Not all species have all the subunits of the Dam1-C. Hence, to avoid noise due to lack of information, only if a species had two or more subunits in each arm, a species was added to the concatenated alignment. The sequences of the subunits were aligned using MAFFT E-INS-i (MAFFT v7.271). To align the structural paralogs MAFFT merge E-INS-i was used. By using MAFFT merge the sub-multiple sequence alignment (MSA) is preserved. The MSA was filtered using trimAL gt 0.05 (trimAl v1.4.rev15). IQ-TREE was used as describe above, the model used was LG+F+R5.

For the tree based on the eight paralogous subunits (Figure4B, all alignments of the subunits were concatenated filtered as previously described, and IQtree was run with model finder and used the LG+F+R5 model

### Topology testing

To test if the tree from the arm based concatenated phylogenetic tree is any different from what we believe is the root of the eukaryotic tree, a topology test was performed. First, trees were built constraining them based on the split in the major supergroups. The arms of the tree were simultaneously constrained (supplementary figure 5). IQtree was used to do topology testing to assess if the constrained trees have a substantially lower likelihood the unconstrained tree. The model used was LG+F+R5.

## Supporting information

Fasta files

## Supplementary Material

Supplementary figures

Supplementary files

## Data Availability Statements

The data underlying this article are available in the article and in its online supplementary material.

## Acknowledgements

The authors thank Leny van Wijk, John van Dam and Eva Deutekom for providing the eukaryotic proteome set. This work is part of the VICI research program with project number 016.160.638 of the Netherlands Organisation for Scientific Research (NWO).

