## Supplementary Figures for "Increased sampling and intra-complex homologies favor vertical over horizontal inheritance of the Dam1 complex"

### Supplementary Material

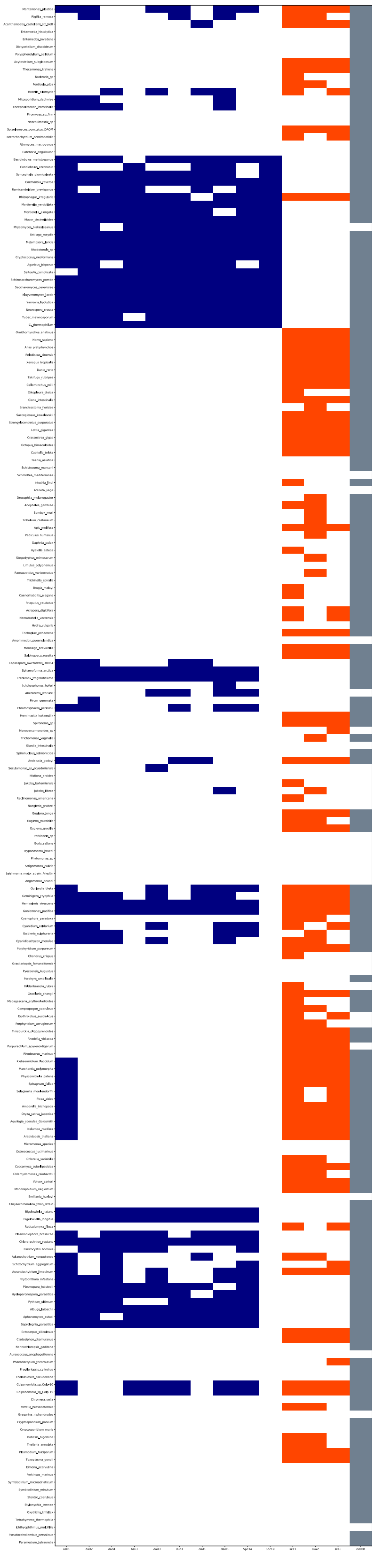

*Figure S1. Full present absent  table Dam1-C subunits, Ska-C subunits and NDC80. Related to figure 2. The presents of all subunits of  Dam1-C(blue), Ska-C (orange)and NDC80(grey).*

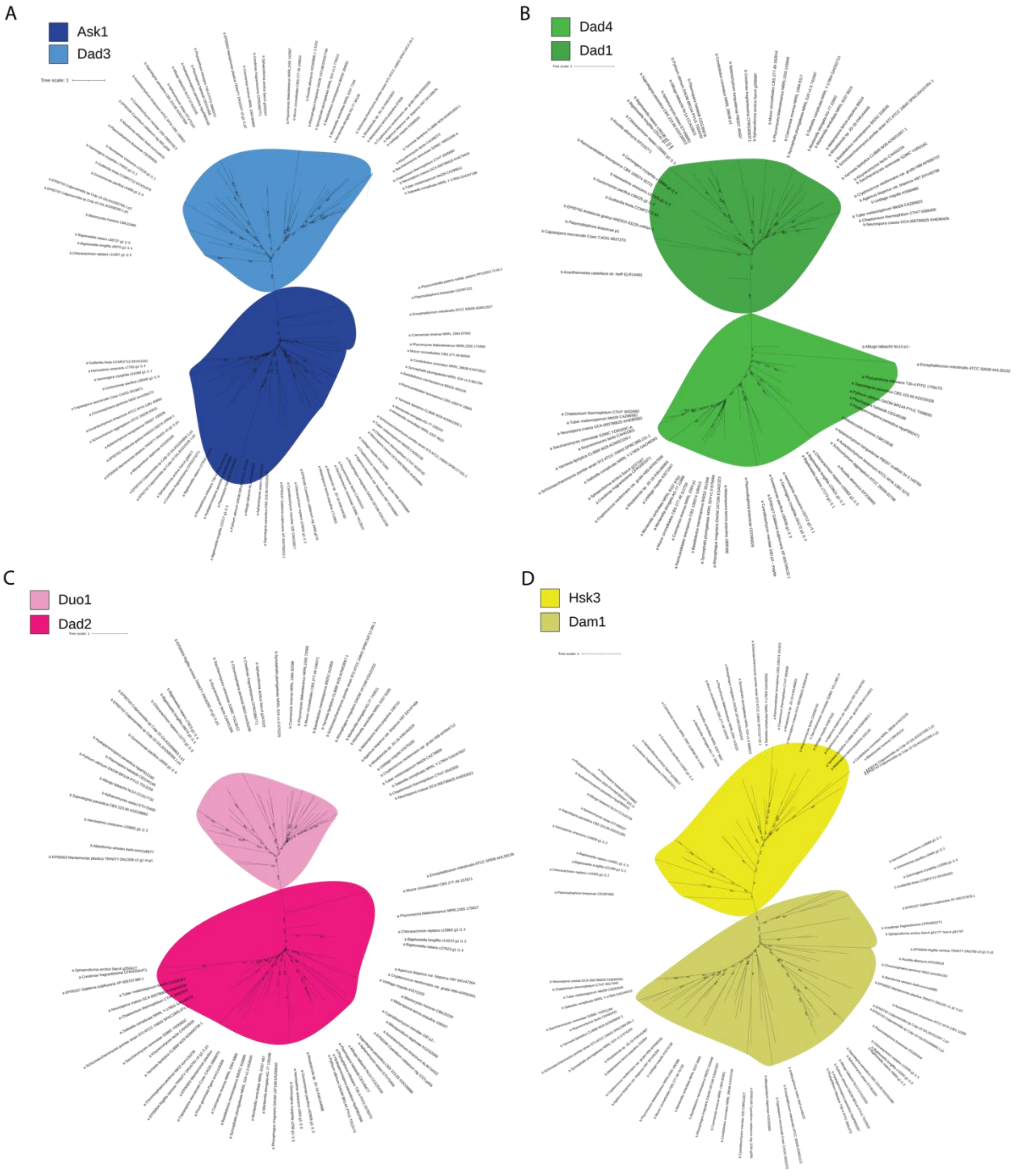


*Figure 2 Gene trees for Dam1-C paralogues pairs. The leaves contain the identifiers of the protein sequences. A) Subunits tree of Ask1(a) and Dad3(b) Random seed number: 473931, Model of substitution: VT+F+R4, B) Subunits tree Dad1(a) and Dad4(b) Random seed number: 804262, Model of substitution: LG+R5. C) Subunit tree Duo1(a) and Dad2(b) Random seed number:87643, Model of substitution VT+F+R4. D) Subunits tree Hsk3(a) and Dam1(b) Random seed number: 695565 Model of substitution LG+F+R5.*

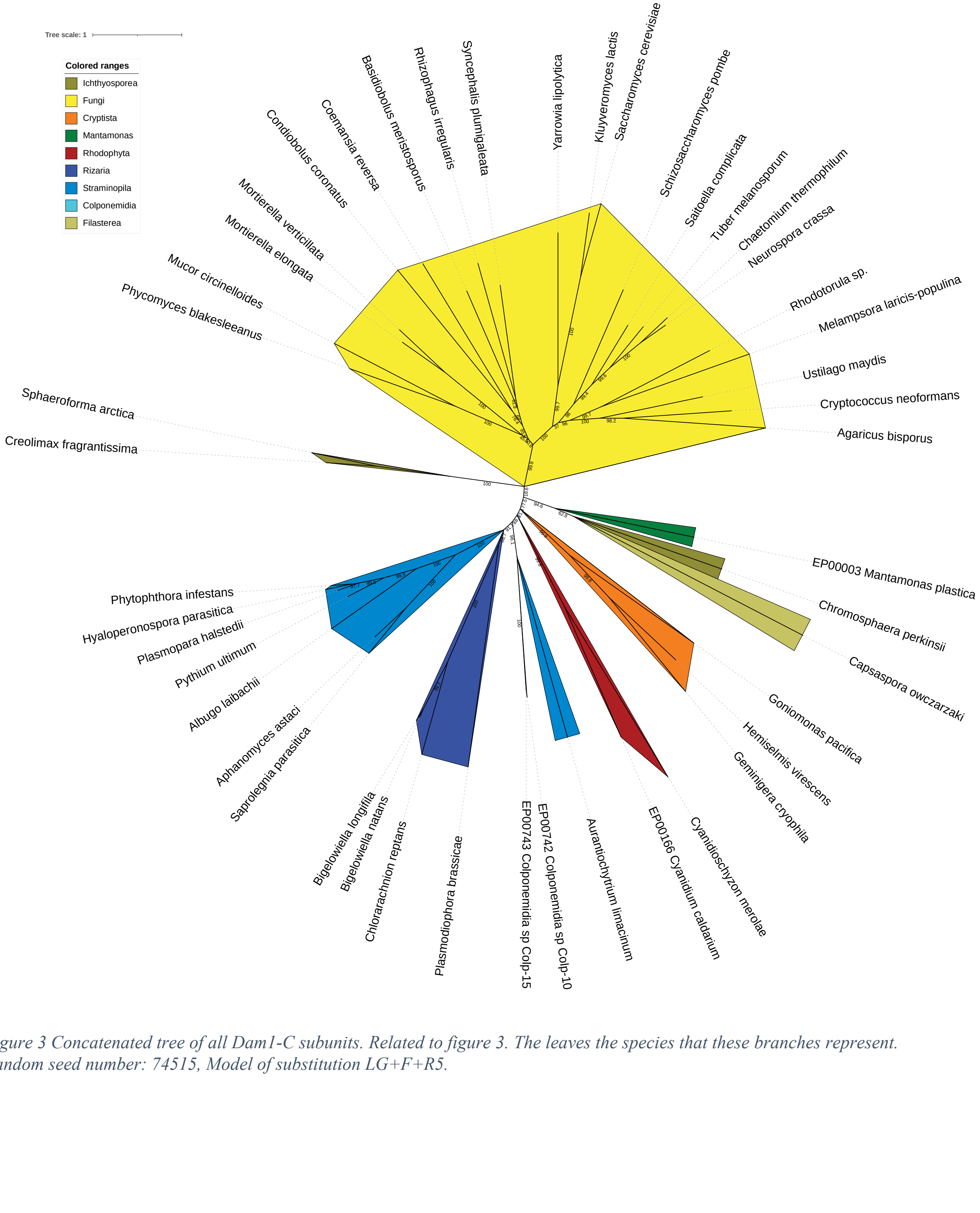


*Figure 3 Concatenated tree of all Dam1-C subunits. Related to figure 3. The leaves the species that these branches represent. Random seed number: 74515, Model of substitution LG+F+R5.*

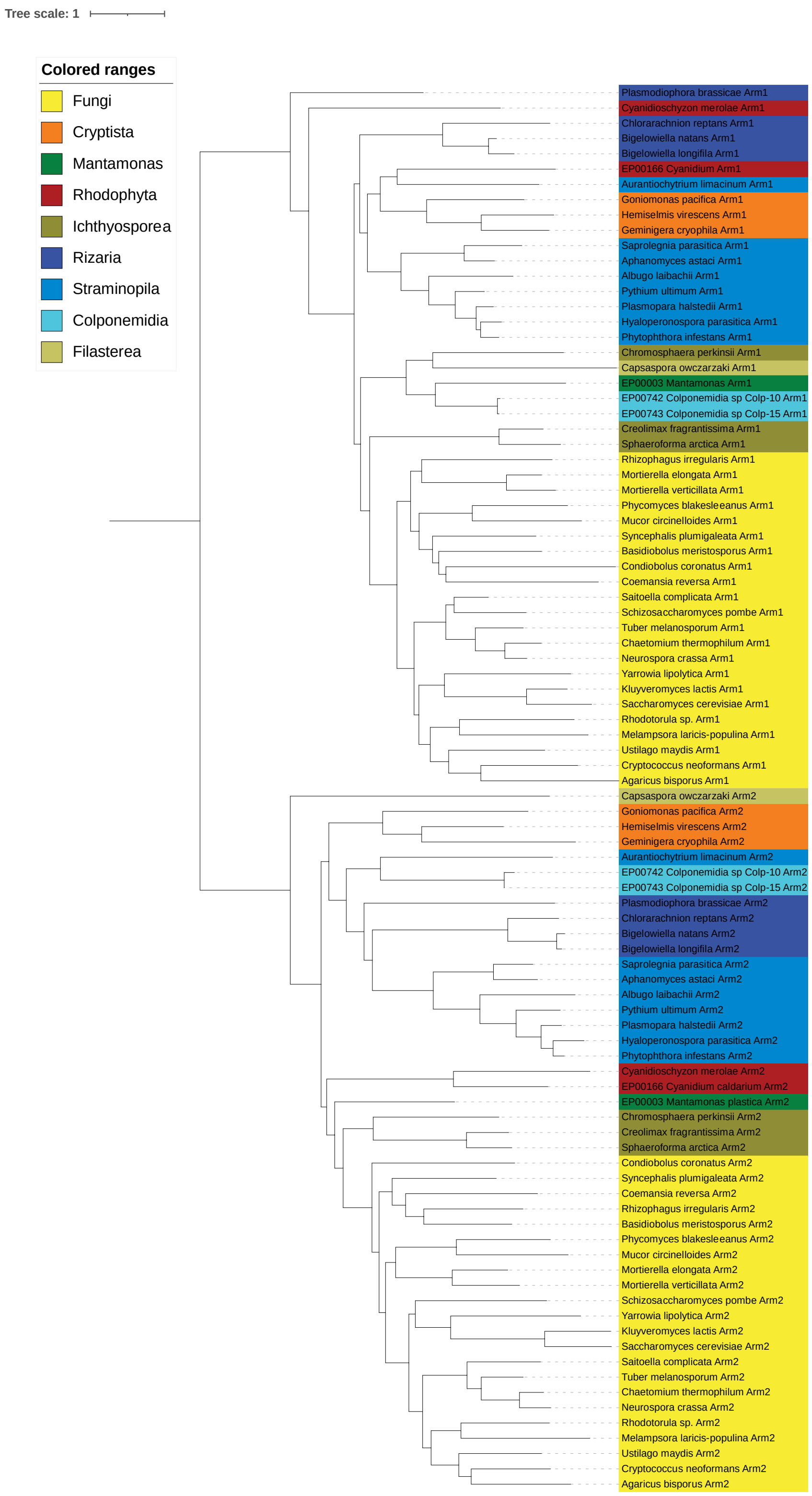


*Figure 4 Phylogeny inferred from aligning the subunits pairs. Related to figure 3. The leaves the species that these branches represent. Random seed number: 691067, Model of substitution LG+F+R5.*

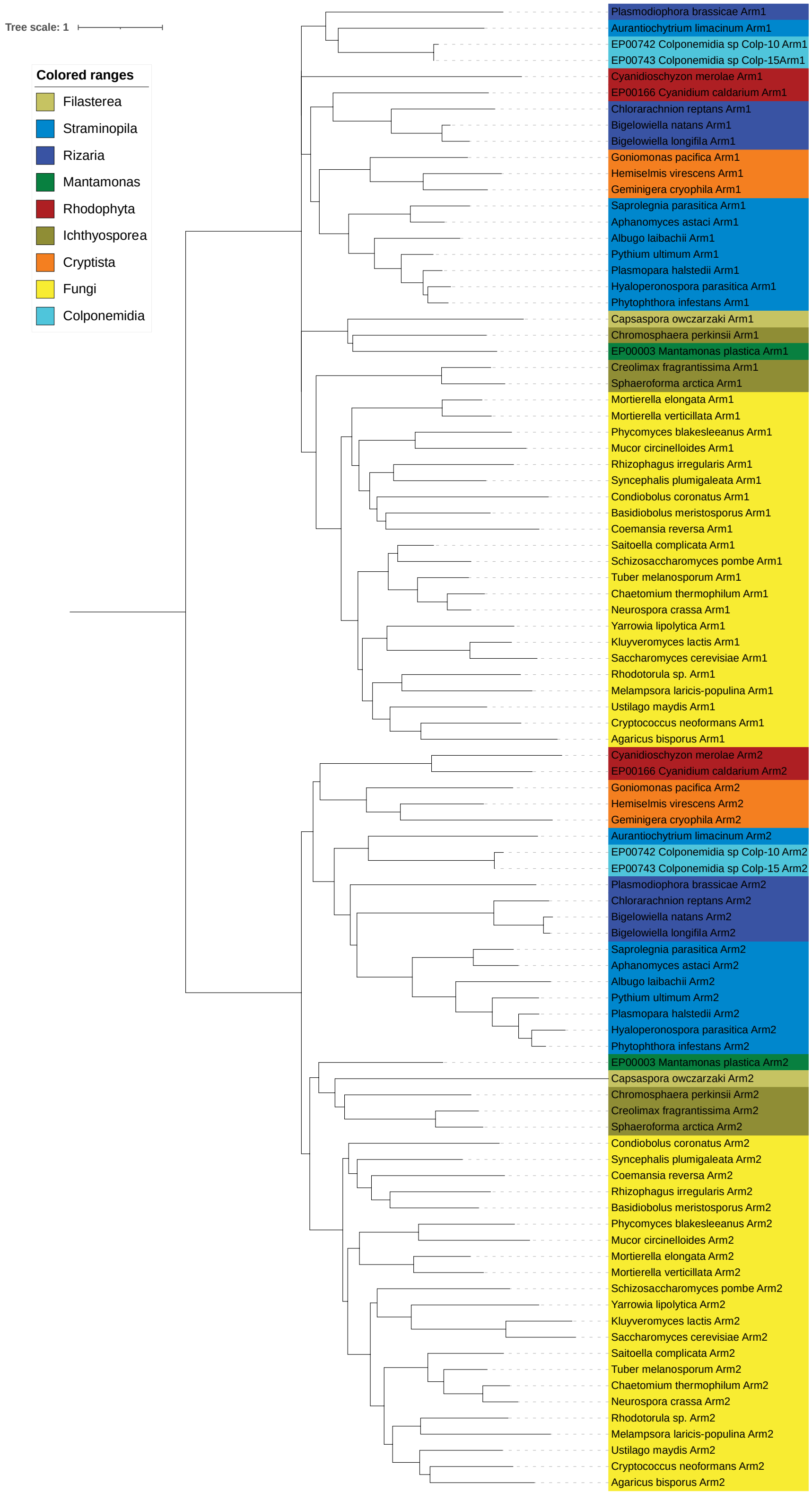
Tree scale: 1
Colored ranges
Filasterea
Straminopila
Rizaria
Mantamonas
Rhodophyta
Ichthyosporea
Cryptista
Fungi
Colponemidia
Plasmodiophora brassicae Arm1
Aurantiochytrium limacinum Arm1
EP00742 Colponemidia sp Colp-10 Arm1
EP00743 Colponemidia sp Colp-15Arm1
Cyanidioschyzon merolae Arm1
EP00166 Cyanidium caldarium Arm1
Chlorarachnion reptans Arm1
Bigelowiella natans Arm1
Bigelowiella longifila Arm1
Goniomonas pacifica Arm1
Hemiselmis virescens Arm1
Geminigera cryophila Arm1
Saprolegnia parasitica Arm1
Aphanomyces astaci Arm1
Albugo laibachii Arm1
Pythium ultimum Arm1
Plasmopara halstedii Arm1
Hyaloperonospora parasitica Arm1
Phytophthora infestans Arm1
Capsaspora owczarzaki Arm1
Chromosphaera perkinsii Arm1
EP00003 Mantamonas plastica Arm1
Creolimax fragrantissima Arm1
Sphaeroforma arctica Arm1
Mortierella elongata Arm1
Mortierella verticillata Arm1
Phycomyces blakesleeanus Arm1
Mucor circinelloides Arm1
Rhizophagus irregularis Arm1
Syncephalis plumigaleata Arm1
Condiobolus coronatus Arm1
Basidiobolus meristosporus Arm1
Coemansia reversa Arm1
Saitoella complicata Arm1
Schizosaccharomyces pombe Arm1
Tuber melanosporum Arm1
Chaetomium thermophilum Arm1
Neurospora crassa Arm1
Yarrowia lipolytica Arm1
Kluyveromyces lactis Arm1
Saccharomyces cerevisiae Arm1
Rhodotorula sp. Arm1
Melampsora laricis-populina Arm1
Ustilago maydis Arm1
Cryptococcus neoformans Arm1
Agaricus bisporus Arm1
Cyanidioschyzon merolae Arm2
EP00166 Cyanidium caldarium Arm2
Goniomonas pacifica Arm2
Hemiselmis virescens Arm2
Geminigera cryophila Arm2
Aurantiochytrium limacinum Arm2
EP00742 Colponemidia sp Colp-10 Arm2
EP00743 Colponemidia sp Colp-15 Arm2
Plasmodiophora brassicae Arm2
Chlorarachnion reptans Arm2
Bigelowiella natans Arm2
Bigelowiella longifila Arm2
Saprolegnia parasitica Arm2
Aphanomyces astaci Arm2
Albugo laibachii Arm2
Pythium ultimum Arm2
Plasmopara halstedii Arm2
Hyaloperonospora parasitica Arm2
Phytophthora infestans Arm2
EP00003 Mantamonas plastica Arm2
Capsaspora owczarzaki Arm2
Chromosphaera perkinsii Arm2
Creolimax fragrantissima Arm2
Sphaeroforma arctica Arm2
Condiobolus coronatus Arm2
Syncephalis plumigaleata Arm2
Coemansia reversa Arm2
Rhizophagus irregularis Arm2
Basidiobolus meristosporus Arm2
Phycomyces blakesleeanus Arm2
Mucor circinelloides Arm2
Mortierella elongata Arm2
Mortierella verticillata Arm2
Schizosaccharomyces pombe Arm2
Yarrowia lipolytica Arm2
Kluyveromyces lactis Arm2
Saccharomyces cerevisiae Arm2
Saitoella complicata Arm2
Tuber melanosporum Arm2
Chaetomium thermophilum Arm2
Neurospora crassa Arm2
Rhodotorula sp. Arm2
Melampsora laricis-populina Arm2
Ustilago maydis Arm2
Cryptococcus neoformans Arm2
Agaricus bisporus Arm2

S

*Figure 5 The constrained phylogeny for  topology testing. Related to figure 3.  Random seed number: 99061, Model of substitution LG+F+R5.*

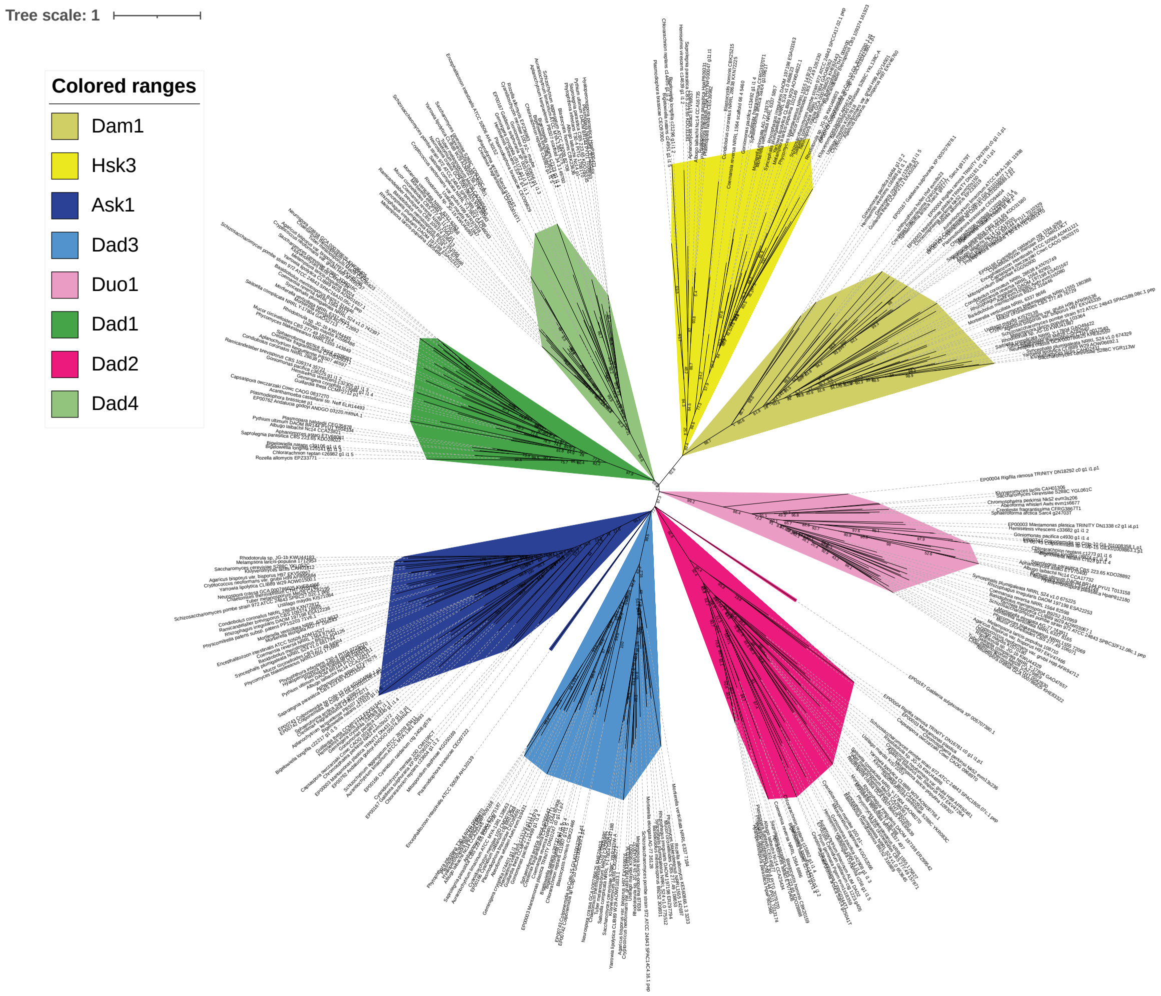


*Figure 6 Phylogeny if the 8 paralogous Dam1-C subunits.  Related to figure 4. The leaves contain the species namen and protein identifier.  Random seed number: 333847, Model of substitution: VT+ R6*

*Table S1 Sources of proteomes*

| Taxonomy ID | Scientific name | genome/ transcirptome | Data Source | Data Source Name | Download date |
|---|---|---|---|---|---|
| **5755** | Acanthamoeba castellanii str Neff | genome | ftp://ftp.ensemblgenomes.org/pub/protists/release-34/fasta/protists_amoebozoa1_collection/acanthamoeba_castellanii_str_neff/pep/Acanthamoeba_castellanii_str_neff.Acastellanii_strNEFF_v1.pep.all.fa.gz | Acastellanii_strNEFF_v1 | 20170124 |
| **5759** | Entamoeba histolytica | genome | ftp://ftp.ensemblgenomes.org/pub/protists/release-34/fasta/entamoeba_histolytica/pep/Entamoeba_histolytica.JCVI-ESG2-1.0.pep.all.fa.gz | JCVI-ESG2-1.0 | 20170124 |
| **370355** | Entamoeba invadens | genome | ftp://ftp.ensemblgenomes.org/pub/protists/release-34/fasta/protists_amoebozoa1_collection/entamoeba_invadens_ip1/pep/Entamoeba_invadens_ip1.EIA2_v2.pep.all.fa.gz | EIA2_v2 | 20170124 |
| **44689** | Dictyostelium discoideum | genome | ftp://ftp.ensemblgenomes.org/pub/protists/release-34/fasta/dictyostelium_discoideum/pep/Dictyostelium_discoideum.dicty_2.7.pep.all.fa.gz | dicty_2.7 | 20170124 |
| **13642** | Polysphondylium pallidum | genome | ftp://ftp.ensemblgenomes.org/pub/protists/release-34/fasta/protists_amoebozoa1_collection/polysphondylium_pallidum_pn500/pep/Polysphondylium_pallidum_pn500.PolPal_Dec2009.pep.all.fa.gz | PolPal_Dec2009 | 20170124 |
| **361139** | Acytostelium subglobosum | genome | ftp://ftp.ncbi.nlm.nih.gov/genomes/all/GCF/000/787/575/GCF_000787575.1_Asub_2.0/GCF_000787575.1_Asub_2.0_protein.faa.gz | Asub_2.0 | 20170124 |
| **3055** | Chlamydomonas reinhardtii | genome | ftp://ftp.ensemblgenomes.org/pub/plants/release-34/fasta/chlamydomonas_reinhardtii/pep/Chlamydomonas_reinhardtii.v3.1.pep.all.fa.gz | v3.1 | 20170124 |
| **145388** | Monoraphidium neglectum | genome | ftp://ftp.ncbi.nlm.nih.gov/genomes/all/GCF/000/611/645/GCF_000611645.1_mono_v1/GCF_000611645.1_mono_v1_protein.faa.gz | GCF_000611645.1 | 20170124 |
| **3067** | Volvox carteri | genome | http://genome.jgi.doe.gov/pages/dynamicOrganismDownload.jsf?organism=Phytozome | V2.1 | 20170124 |
| **3175** | Klebsormidium flaccidum | genome | http://genome.jgi.doe.gov/pages/dynamicOrganismDownload.jsf?organism=Phytozome | | 20170124 |
| **296587** | Micromonas species | genome | https://phytozome.jgi.doe.gov/biomart/martview/739c2860d5e06348b925fe0f410cb17c filename: Micromonas_species_RCC299_mart_export.txt | | 20170124 |
| **242159** | Ostreococcus lucimarinus | genome | ftp://ftp.ensemblgenomes.org/pub/plants/release-34/fasta/ostreococcus_lucimarinus/pep/Ostreococcus_lucimarinus.ASM9206v1.pep.all.fa.gz | ASM9206v1 | 20170124 |
| **554065** | Chlorella variabilis | genome | http://genome.jgi.doe.gov/ChlNC64A_1/download/Chlorella_NC64A.best_proteins.fasta.gz | | 20170117 |
| **248742** | Coccomyxa subellipsoidea | genome | https://phytozome.jgi.doe.gov/biomart/martview/739c2860d5e06348b925fe0f410cb17c filename: Coccomyxa_subellipsoidea_mart_export.txt | | 20170124 |
| **3329** | Picea abies | genome | ftp://plantgenie.org/Data/ConGenIE/Picea_abies/v1.0/FASTA/GenePrediction/Pabies1.01.0-HC-pep.faa.gz, | v1.0 | 20170124 |

| | | | | | |
|---|---|---|---|---|---|
| | | | ftp://plantgenie.org/Data/ConGenIE/Picea_abies/v1.0/FASTA/GenePrediction/Pabies1.01.0-MC-pep.faa.gz | | |
| **3197** | Marchantia polymorpha | genome | https://phytozome.jgi.doe.gov/biomart/martview/739c2860d5e06348b925fe0f410cb17c filename: Marchantia_polymorpha_mart_export.txt | | 20170124 |
| **3218** | Physcomitrella patens | genome | ftp://ftp.ensemblgenomes.org/pub/plants/release-34/fasta/physcomitrella_patens/pep/Physcomitrella_patens.ASM242v1.pep.all.fa.gz | ASM242v1 | 20170124 |
| **53036** | Sphagnum fallax | genome | https://phytozome.jgi.doe.gov/biomart/martview/739c2860d5e06348b925fe0f410cb17c filename: Sphagnum_fallax_mart_export.txt | | 20170124 |
| **4432** | Nelumbo nucifera | genome | ftp://ftp.ncbi.nlm.nih.gov/genomes/all/GCF/000/365/185/GCF_000365185.1_Chinese_Lotus_1.1/ | GCF_000365185.1 | 20170327 |
| **3702** | Arabidopsis thaliana | genome | ftp://ftp.ensemblgenomes.org/pub/plants/release-34/fasta/arabidopsis_thaliana/pep/Arabidopsis_thaliana.TAIR10.pep.all.fa.gz | TAIR10 | 20170124 |
| **218851** | Aquilegia coerulea Goldsmith | genome | https://phytozome.jgi.doe.gov/biomart/martview/739c2860d5e06348b925fe0f410cb17c filename: Aquilegia coerulea_mart_export.txt | | 20170124 |
| **13333** | Amborella trichopoda | genome | ftp://ftp.ensemblgenomes.org/pub/plants/release-34/fasta/amborella_trichopoda/pep/Amborella_trichopoda.AMTR1.0.pep.all.fa.gz | AMTR1.0 | 20170124 |
| **39947** | Oryza sativa japonica | genome | ftp://ftp.ensemblgenomes.org/pub/plants/release-34/fasta/oryza_sativa/pep/Oryza_sativa.IRGSP-1.0.pep.all.fa.gz | IRGSP-1.0 | 20170124 |
| **88036** | Selaginella moellendorffii | genome | https://phytozome.jgi.doe.gov/biomart/martview/739c2860d5e06348b925fe0f410cb17c filename: Selaginella_moellendorffii_mart_export.txt | v1.0 | 20170327 |
| **2762** | Cyanophora paradoxa | genome | http://cyanophora.rutgers.edu/cyanophora/Cyanophora_paradoxa_MAKER_gene_predictions-022111-aa.fasta | | 20170124 |
| **130081** | Galdieria sulphuraria | genome | ftp://ftp.ncbi.nlm.nih.gov/genomes/all/GCF/000/341/285/GCF_000341285.1_ASM34128v1/GCF_000341285.1_ASM34128v1_protein.faa.gz | | 20210527 |
| **45157** | Cyanidioschyzon merolae | genome | ftp://ftp.ensemblgenomes.org/pub/release-34/plants/fasta/cyanidioschyzon_merolae/pep/ | ASM9120v1 | 20170117 |
| **35688** | Porphyridium purpureum | genome | http://cyanophora.rutgers.edu/porphyridium/Porphyridium_genemodels_UPDATED.fasta | | 20170124 |
| **2769** | Chondrus crispus | genome | ftp://ftp.ensemblgenomes.org/pub/release-34/plants/fasta/plants_rhodophyta1_collection/chondrus_crispus/pep/ | ASM35022v2 | 20170117 |
| **2782** | Gracilariopsis lemaneiformis | genome | https://ftp.ncbi.nlm.nih.gov/genomes/all/GCA/003/346/895/GCA_003346895.1_Glem_v01/GCA_003346895.1_Glem_v01_genomic.fna.gz | v01 | 20210527 |
| **2771** | Cyanidium caldarium | genome | http://cyanophora.rutgers.edu/redEST/redESTs.tar.gz | Cyanidium.protein | 20210527 |
| **2787** | Porphyra purpurea | transcriptome | https://www.ncbi.nlm.nih.gov/sra/?term=txid2787[Organism:noexp] | Porphyra purpurea EST project | 20210527 |
| **2786** | Porphyra umbilicalis | genome | ftp://ftp.ncbi.nlm.nih.gov/genomes/all/GCA/002/049/455/GCA_002049455.2_P_umbilicalis_v1/GCA_002049455.2_P_umbilicalis_v1_protein.faa.gz | v1 | 20210527 |

| | | | | | |
|---|---|---|---|---|---|
| **2788** | Pyropia yezoensis | genome | http://nrifs.fra.affrc.go.jp/cgi-bin/lime_download/lime.cgi?nori_FASTA_AminoAcid | Ver. 1 | 20210527 |
| **48942** | Calliarthron tuberculosum | genome | http://realdb.algaegenome.org/P.c.t.html# | Calliarthron_tuberculosum_aa | 20210527 |
| **31481** | Hildenbrandia rubra | transcriptome | http://cyanophora.rutgers.edu/redEST/redESTs.tar.gz | est | 20210527 |
| **2822** | Palmaria palmata | transcriptome | http://cyanophora.rutgers.edu/redEST/redESTs.tar.gz | est | 20210527 |
| **291168** | Griffithsia okiensis | EST | https://www.ncbi.nlm.nih.gov/nuccore?LinkName=biosample_nuccore&from_uid=167350 | LIBEST_024364 | 20210527 |
| **28020** | Furcellaria lumbricalis | EST | https://www.ncbi.nlm.nih.gov/nuccore?LinkName=biosample_nuccore&from_uid=169272,https://www.ncbi.nlm.nih.gov/nuccore?LinkName=biosample_nuccore&from_uid=169271 | LIBEST_026199,LIBEST_026198 | 20210527 |
| **2769** | Chondrus crispus | genome | ftp://ftp.ncbi.nlm.nih.gov/genomes/all/GCF/000/350/225/GCF_000350225.1_ASM35022v2/GCF_000350225.1_ASM35022v2_protein.faa.gz | ASM35022v2 | 20210527 |
| **305493** | Eucheuma denticulatum | transcriptome | https://www.ncbi.nlm.nih.gov/sites/nuccore?term=379657[BioProject] | GFKZ01000000 | 20210527 |
| **172969** | Gracilaria changii | transcriptome | https://www.ncbi.nlm.nih.gov/sra?LinkName=biosample_sra&from_uid=7306656,https://www.ncbi.nlm.nih.gov/sra?LinkName=biosample_sra&from_uid=7306657 | GC_UT_1,GC_UT_2 | 20210527 |
| **753684** | Madagascaria erythrocladioides | transcriptome | https://doi.org/10.6084/m9.figshare.12410606 | MMETSP1450 | 20210527 |
| **867925** | Rhodochaete pulchella | transcriptome | http://cyanophora.rutgers.edu/redEST/redESTs.tar.gz | est | 20210527 |
| **31354** | Compsopogon caeruleus | transcriptome | https://doi.org/10.6084/m9.figshare.12410606 | MMETSP0312 | 20210527 |
| **1077150** | Erythrolobus australicus | transcriptome | https://doi.org/10.6084/m9.figshare.12410606 | MMETSP1353 | 20210527 |
| **708628** | Erythrolobus madagascarensis | transcriptome | https://doi.org/10.6084/m9.figshare.12410606 | MMETSP1354 | 20210527 |
| **2792** | Porphyridium aerugineum | transcriptome | https://doi.org/10.6084/m9.figshare.12410606 | MMETSP0313 | 20210527 |
| **35688** | Porphyridium purpureum | genome | http://porphyra.rutgers.edu/Porphyridium_purpureum_v2_genome_data.zip | v.2 | 20210527 |
| **708627** | Timspurckia oligopyrenoides | transcriptome | https://doi.org/10.6084/m9.figshare.12410606 | MMETSP1172 | 20210527 |
| **2801** | Rhodella violacea | transcriptome | https://doi.org/10.6084/m9.figshare.12410606 | MMETSP0167,MMETSP0314 | 20210527 |
| **282340** | Purpureofilum apyrenoidigerum | transcriptome | http://cyanophora.rutgers.edu/redEST/redESTs.tar.gz | est | 20210527 |
| **101924** | Rhodosorus marinus | transcriptome | https://doi.org/10.6084/m9.figshare.12410606 | MMETSP0011,MMETSP0315 | 20210527 |
| **446134** | Stylonematophyceae sp CCMP1999 | transcriptome | https://doi.org/10.6084/m9.figshare.12410606 | MMETSP1475 | 20210527 |
| **46947** | Geminigera cryophila | genome | https://figshare.com/articles/Marine_Microbial_Eukaryotic_Transcriptome_Sequencing_Project_re-assemblies/3840153/3 | | 20201020 |
| **77927** | Hemiselmis virescens | genome | https://figshare.com/articles/Marine_Microbial_Eukaryotic_Transcriptome_Sequencing_Project_re-assemblies/3840153/3 | | 20201020 |
| **55529** | Guillardia theta | genome | ftp://ftp.ensemblgenomes.org/pub/protists/release-34/fasta/guillardia_theta/pep/ | GCA_000315625.1 | 20170117 |

| | | | | | |
|---|---|---|---|---|---|
| **195067** | Goniomonas pacifica | genome | https://figshare.com/articles/Marine_Microbial_Eukaryotic_Transcriptome_Sequencing_Project_re-assemblies/3840153/3 | | 20210527 |
| **856889** | Mantamonas plastica | transcriptome | https://trace.ncbi.nlm.nih.gov/Traces/sra/?run=SRR5997433 | Mantamonas plastica CCAP 1946/1 transcriptome | 20210527 |
| **1122280** | Rigifila ramosa | transcriptome | https://trace.ncbi.nlm.nih.gov/Traces/sra/?run=SRR5997435 | Rigifila ramosa CCAP 1967/1 transcriptome | 20210527 |
| **3039** | Euglena gracilis | genome | ftp://ftp.pride.ebi.ac.uk/pride/data/archive/2019/01/PXD009998/Euglena_translated_transcriptome.fasta | translated_transcriptome | 20210527 |
| **3037** | Euglena longa | transcriptome | ftp://ftp.ncbi.nlm.nih.gov/sra/wgs_aux/GG/OE/GGOE01/GGOE01.1.fsa_nt.gz | GGOE01000000 | 20210527 |
| **38275** | Euglena mutabilis | transcriptome | https://trace.ncbi.nlm.nih.gov/Traces/sra/?run=ERR351290,https://trace.ncbi.nlm.nih.gov/Traces/sra/?run=ERR351289,https://trace.ncbi.nlm.nih.gov/Traces/sra/?run=ERR351288,https://trace.ncbi.nlm.nih.gov/Traces/sra/?run=ERR351287,https://trace.ncbi.nlm.nih.gov/Traces/sra/?run=ERR351286,https://trace.ncbi.nlm.nih.gov/Traces/sra/?run=ERR351285 | Population Genomics of Euglena mutabilis | 20210527 |
| **75058** | Bodo saltans | genome | ftp://ftp.ensemblgenomes.org/pub/release-34/protists/fasta/protists_euglenozoa1_collection/bodo_saltans/pep/ | | 20170105 |
| **1314962** | Perkinsela sp | genome | ftp://ftp.ensemblgenomes.org/pub/release-34/protists/fasta/protists_euglenozoa1_collection/perkinsela_sp_ccap_1560_4/pep/ | ASM123584v1 | 20170105 |
| **347515** | Leishmania major strain Friedlin | genome | ftp://ftp.ensemblgenomes.org/pub/release-34/protists/fasta/leishmania_major/pep/ | ASM272v2 | 20170105 |
| **5691** | Trypanosoma brucei | genome | ftp://ftp.ensemblgenomes.org/pub/release-34/protists/fasta/trypanosoma_brucei/pep/ | Chr11 | 20170105 |
| **59799** | Angomonas deanei | genome | ftp://ftp.ensemblgenomes.org/pub/release-34/protists/fasta/protists_euglenozoa1_collection/angomonas_deanei/pep/ | GCA_000442575.2 | 20170105 |
| **28005** | Strigomonas culicis | genome | ftp://ftp.ensemblgenomes.org/pub/release-34/protists/fasta/protists_euglenozoa1_collection/strigomonas_culicis/pep/ | GCA_000442495.1 | 20170105 |
| **134013** | Phytomonas sp | genome | ftp://ftp.ensemblgenomes.org/pub/release-34/protists/fasta/protists_euglenozoa1_collection/phytomonas_sp_isolate_hart1/pep/ | AKI_PRJEB1539_v1 | 20170105 |
| **5741** | Giardia intestinalis | genome | http://giardiadb.org/common/downloads/release-29/GintestinalisAssemblageAWB/fasta/data/ | GiardiaDB-29 Assemblage A isolate WB | 20170105 |
| **348837** | Spironucleus salmonicida | genome | ftp://ftp.ensemblgenomes.org/pub/release-34/protists/fasta/protists_fornicata1_collection/spironucleus_salmonicida/pep/ | SSK3.0 | 20170105 |
| **453998** | Monocercomonoides sp | genome | http://www.protistologie.cz/hampllab/data.html | Mono14B | 20170303 |

| | | | | | |
|---|---|---|---|---|---|
| **5722** | Trichomonas vaginalis | genome | ftp://ftp.ensemblgenomes.org/pub/release-34/protists/fasta/protists_parabasalia1_collection/trichomonas_vaginalis_g3/pep/ | G3 | 20170105 |
| **5762** | Naegleria gruberi | genome | ftp://ftp.ensemblgenomes.org/pub/release-34/protists/fasta/protists_heterolobosea1_collection/naegleria_gruberi/pep/ | v1 | 20170105 |
| **2903** | Emiliania huxleyi | genome | ftp://ftp.ensemblgenomes.org/pub/protists/release-34/fasta/emiliania_huxleyi/pep/ | GCA_000372725.1 | 20170117 |
| **1460289** | Chrysochromulina tobin strain | genome | ftp://ftp.ncbi.nlm.nih.gov/genomes/all/GCA/001/275/005/GCA_001275005.1_Ctobinv2/GCA_001275005.1_Ctobinv2_protein.faa.gz | GCA_001275005.1 v2 | 20170117 |
| **2027451** | Hemimastix kukwesjijk | transcriptome | https://datadryad.org/resource/doi:10.5061/dryad.n5g39d7/4 | assembly_July2016 | 20181121 |
| **2027454** | Spironema sp | transcriptome | https://datadryad.org/resource/doi:10.5061/dryad.n5g39d7/4 | assembly_July2016 | 20181121 |
| **7955** | Danio rerio | genome | ftp://ftp.ensembl.org/pub/release-87/fasta/danio_rerio/pep/Danio_rerio.GRCz10.pep.all.fa.gz | GRCz10 | 20170119 |
| **31033** | Takifugu rubripes | genome | ftp://ftp.ncbi.nlm.nih.gov/genomes/all/GCF/000/180/615/GCF_000180615.1_FUGU5/GCF_000180615.1_FUGU5_protein.faa.gz | FUGU5 | 20170119 |
| **8364** | Xenopus tropicalis | genome | ftp://ftp.ensembl.org/pub/release-87/fasta/xenopus_tropicalis/pep/ | Xenopus_tropicalis_JGI_4.2 | 20170119 |
| **294128** | Hyalella azteca | genome | ftp://ftp.ncbi.nlm.nih.gov/genomes/all/GCF/000/764/305/GCF_000764305.1_Hazt_2.0/GCF_000764305.1_Hazt_2.0_protein.faa.gz | Hazt_2.0 | 20170119 |
| **283909** | Capitella teleta | genome | ftp://ftp.ensemblgenomes.org/pub/metazoa/release-34/fasta/capitella_teleta/pep/Capitella_teleta.GCA_000328365.1.pep.all.fa.gz | Capitella teleta v1.0 | 20170119 |
| **407821** | Stegodyphus mimosarum | genome | ftp://ftp.ensemblgenomes.org/pub/metazoa/release-34/fasta/stegodyphus_mimosarum/pep/Stegodyphus_mimosarum.GCA_000611955.2.pep.all.fa.gz | Stegodyphus_mimosarum_v1 | 20170119 |
| **8839** | Anas platyrhynchos | genome | ftp://ftp.ensembl.org/pub/release-87/fasta/anas_platyrhynchos/pep/Anas_platyrhynchos.BGI_duck_1.0.pep.all.fa.gz | BGI_duck_1.0 | 20170119 |
| **29159** | Crassostrea gigas | genome | ftp://ftp.ensemblgenomes.org/pub/metazoa/release-34/fasta/crassostrea_gigas/pep/Crassostrea_gigas.GCA_000297895.1.pep.all.fa.gz | oyster_v9 | 20170119 |
| **6669** | Daphnia pulex | genome | ftp://ftp.ensemblgenomes.org/pub/metazoa/release-34/fasta/daphnia_pulex/pep/Daphnia_pulex.GCA_000187875.1.pep.all.fa.gz | V1.0 | 20170119 |
| **7739** | Branchiostoma floridae | genome | ftp://ftp.ncbi.nlm.nih.gov/genomes/all/GCA/000/003/815/GCA_000003815.1_Version_2/GCA_000003815.1_Version_2_protein.faa.gz | GCA_000003815.1_Version_2_genomic | 20170119 |

| | | | | | |
|---|---|---|---|---|---|
| **37653** | Octopus bimaculoides | genome | ftp://ftp.ensemblgenomes.org/pub/metazoa/release-34/fasta/octopus_bimaculoides/pep/Octopus_bimaculoides.PRJNA270931.pep.all.fa.gz | Octopus_bimaculoides_v2_0 | 20170119 |
| **7868** | Callorhinchus milii | genome | ftp://ftp.ncbi.nlm.nih.gov/genomes/all/GCF/000/165/045/GCF_000165045.1_Callorhinchus_milii-6.1.3/GCF_000165045.1_Callorhinchus_milii-6.1.3_protein.faa.gz | Callorhinchus_milii-6.1.3 | 20170119 |
| **70779** | Acropora digitifera | genome | ftp://ftp.ncbi.nlm.nih.gov/genomes/all/GCF/000/222/465/GCF_000222465.1_Adig_1.1/GCF_000222465.1_Adig_1.1_protein.faa.gz | Adig1.1 | 20170119 |
| **45351** | Nematostella vectensis | genome | ftp://ftp.ensemblgenomes.org/pub/metazoa/release-34/fasta/nematostella_vectensis/pep/Nematostella_vectensis.GCA_000209225.1.pep.all.fa.gz | ASM20922v1 - NemVe_1 | 20170119 |
| **6087** | Hydra vulgaris | genome | ftp://ftp.ncbi.nlm.nih.gov/genomes/all/GCF/000/004/095/GCF_000004095.1_Hydra_RP_1.0/GCF_000004095.1_Hydra_RP_1.0_protein.faa.gz | Hydra_RP-1.0 | 20170119 |
| **7070** | Tribolium castaneum | genome | ftp://ftp.ncbi.nlm.nih.gov/genomes/all/GCA/000/002/335/GCA_000002335.3_Tcas5.2/GCA_000002335.3_Tcas5.2_protein.faa.gz | Tcas5.2 | 20170119 |
| **7227** | Drosophila melanogaster | genome | ftp://ftp.ensemblgenomes.org/pub/metazoa/release-34/fasta/drosophila_melanogaster/pep/Drosophila_melanogaster.BDGP6.pep.all.fa.gz | BDGP6 | 20170119 |
| **7165** | Anopheles gambiae | genome | ftp://ftp.ensemblgenomes.org/pub/metazoa/release-34/fasta/anopheles_gambiae/pep/ | Agam4.4 | 20170119 |
| **7668** | Strongylocentrotus purpuratus | genome | http://www.echinobase.org/Echinobase/SpDownloads Version 4.2 | Spur4.2 | 20170119 |
| **225164** | Lottia gigantea | genome | ftp://ftp.ensemblgenomes.org/pub/metazoa/release-34/fasta/lottia_gigantea/pep/Lottia_gigantea.GCA_000327385.1.pep.all.fa.gz | Lotgi1 | 20170119 |
| **10224** | Saccoglossus kowalevskii | genome | ftp://ftp.ncbi.nlm.nih.gov/genomes/all/GCF/000/003/605/GCF_000003605.2_Skow_1.1/GCF_000003605.2_Skow_1.1_protein.faa.gz | Skow_1.1 | 20170119 |
| **7460** | Apis melifera | genome | ftp://ftp.ensemblgenomes.org/pub/metazoa/release-34/fasta/apis_mellifera/pep/Apis_mellifera.GCA_000002195.1.pep.all.fa.gz | Amel_4.5 | 20170119 |
| **7091** | Bombyx mori | genome | ftp://ftp.ensemblgenomes.org/pub/metazoa/release-34/fasta/bombyx_mori/pep/Bombyx_mori.GCA_000151625.1.pep.all.fa.gz | ASM15162v1 | 20170119 |
| **9258** | Ornithorhynchus anatinus | genome | ftp://ftp.ensembl.org/pub/release-87/fasta/ornithorhynchus_anatinus/pep/Ornithorhynchus_anatinus.OANA5.pep.all.fa.gz | OANA5 | 20170119 |
| **9606** | Homo sapiens | genome | ftp://ftp.ensembl.org/pub/release-87/fasta/homo_sapiens/pep/ | GRCH38.p7 | 20170119 |
| **6850** | Limulus polyphemus | genome | ftp://ftp.ncbi.nlm.nih.gov/genomes/all/GCF/000/517/525/GCF_000517525.1_Limulus_polyphemus- | Limulus polyphemus-2.1.2 | 20170119 |

| | | | | | |
|---|---|---|---|---|---|
| | | | 2.1.2/GCF_000517525.1_Limulus_polyphemus-2.1.2_protein.faa.gz | | |
| **1819745** | Intoshia linei | genome | ftp://ftp.ncbi.nlm.nih.gov/genomes/all/GCA/001/642/005/GCA_001642005.1_IntLin1.0/GCA_001642005.1_IntLin1.0_protein.faa.gz | IntLin_1.0 | 20170119 |
| **6279** | Brugia malayi | genome | ftp://ftp.ensemblgenomes.org/pub/metazoa/release-34/fasta/brugia_malayi/pep/Brugia_malayi.B_malayi-3.1.pep.all.fa.gz | B_malayi-3.1 | 20170119 |
| **6239** | Caenorhabditis elegans | genome | ftp://ftp.ensemblgenomes.org/pub/metazoa/release-34/fasta/caenorhabditis_elegans/pep/Caenorhabditis_elegans.WBcel235.pep.all.fa.gz | Wcel235 | 20170119 |
| **6334** | Trichinella spiralis | genome | ftp://ftp.ncbi.nlm.nih.gov/genomes/all/GCF/000/181/795/GCF_000181795.1_Trichinella_spiralis-3.7.1/GCF_000181795.1_Trichinella_spiralis-3.7.1_protein.faa.gz | Trichinella spiralis-3.7.1 | 20170119 |
| **121225** | Pediculus humanus | genome | ftp://ftp.ensemblgenomes.org/pub/metazoa/release-34/fasta/pediculus_humanus/pep/Pediculus_humanus.PhumU2.pep.all.fa.gz | PhumU2 | 20170119 |
| **10228** | Trichoplax adhaerens | genome | ftp://ftp.ensemblgenomes.org/pub/metazoa/release-34/fasta/trichoplax_adhaerens/pep/Trichoplax_adhaerens.ASM15027v1.pep.all.fa.gz | ASM15027v1 | 20170119 |
| **60517** | Taenia asiatica | genome | ftp://ftp.ebi.ac.uk/pub/databases/wormbase/parasite/releases/WBPS8/species/taenia_asiatica/PRJEB532/taenia_asiatica.PRJEB532.WBPS8.protein.fa.gz | T_asiatica_South_Korea_v1_0_4 | 20170119 |
| **79327** | Schmidtea mediterranea | genome | ftp://ftp.ebi.ac.uk/pub/databases/wormbase/parasite/releases/WBPS8/species/schmidtea_mediterranea/PRJNA12585/schmidtea_mediterranea.PRJNA12585.WBPS8.protein.fa.gz or http://smedgd.stowers.org/files/SmedSxl_genome_v4.0.all.maker.proteins.fasta.gz | SmedGD_v1.3 and SmedGD_v4.0 | 20170119 |
| **6183** | Schistosoma mansoni | genome | ftp://ftp.ebi.ac.uk/pub/databases/wormbase/parasite/releases/WBPS8/species/schistosoma_mansoni/PRJEA36577/schistosoma_mansoni.PRJEA36577.WBPS8.protein.fa.gz | ASM23792v2 | 20170119 |
| **400682** | Amphimedon queenslandica | genome | http://amphimedon.qcloud.qcif.edu.au/downloads.html | Aqu2.1 | 20170119 |
| **13735** | Pelodiscus sinensis | genome | ftp://ftp.ensembl.org/pub/release-87/fasta/pelodiscus_sinensis/pep/Pelodiscus_sinensis.PelSin_1.0.pep.all.fa.gz | PelSin_1.0 | 20170119 |
| **104782** | Adineta vaga | genome | http://www.genoscope.cns.fr/adineta/data/Adineta_vaga.v2.pep.fa.gz | Adineta_vaga.v2 | 20170119 |
| **947166** | Ramazzottius varieornatus | genome | ftp://ftp.ncbi.nlm.nih.gov/genomes/all/GCA/001/949/185/GCA_001949185.1_Rvar_4.0/GCA_001949185.1_Rvar_4.0_protein.faa.gz | rvar_4.0 | 20170119 |
| **34765** | Oikopleura dioica | genome | ftp://ftp.ncbi.nlm.nih.gov/genomes/all/GCA/000/209/535/GCA_000209535.1_ASM20953v1/GCA_000209535.1_ASM20953v1_protein.faa.gz | Oikopleura_peptides_reference_v1.0 | 20170119 |

| | | | | | |
|---|---|---|---|---|---|
| **7719** | Ciona intestinalis | genome | ftp://ftp.ensembl.org/pub/release-87/fasta/ciona_intestinalis/pep/Ciona_intestinalis.KH.pep.all.fa.gz | KH | 20170119 |
| **37621** | Priapulus caudatus | genome | ftp://ftp.ncbi.nlm.nih.gov/genomes/all/GCF/000/485/595/GCF_000485595.1_Priapulus_caudatus-5.0.1/GCF_000485595.1_Priapulus_caudatus-5.0.1_protein.faa.gz | Priapulus caudatus-5.0.1 | 20170119 |
| **81824** | Monosiga brevicollis | genome | ftp://ftp.ensemblgenomes.org/pub/protists/release-34/fasta/protists_choanoflagellida1_collection/monosiga_brevicollis_mx1/pep/Monosiga_brevicollis_mx1.V1.0.pep.all.fa.gz | MonBr V1.0, | 20170120 |
| **946362** | Salpingoeca rosetta | genome | ftp://ftp.ensemblgenomes.org/pub/protists/release-34/fasta/protists_choanoflagellida1_collection/salpingoeca_rosetta/pep/Salpingoeca_rosetta.Proterospongia_sp_ATCC50818.pep.all.fa.gz | Proterospongia_sp_ATCC50818 | 20170120 |
| **595528** | Capsaspora owczarzaki 30864 | genome | https://figshare.com/articles/dataset/Genome_-_Capsaspora_owczarzaki_v3_/4123158 | Capsaspora owczarzaki (v3) | 20220920 |
| **5341** | Agaricus bisporus | genome | ftp://ftp.ensemblgenomes.org/pub/fungi/release-34/fasta/fungi_basidiomycota1_collection/agaricus_bisporus_var_bisporus_h97/pep/Agaricus_bisporus_var_bisporus_h97.Agabi_varbisH97_2.pep.all.fa.gz | Agabi_varbisH97_2 | 20170120 |
| **5207** | Cryptococcus neoformans | genome | ftp://ftp.ensemblgenomes.org/pub/fungi/release-34/fasta/fungi_basidiomycota1_collection/cryptococcus_neoformans_var_grubii_h99/pep/Cryptococcus_neoformans_var_grubii_h99.CNA3.pep.all.fa.gz | CNA3 | 20170120 |
| **209285** | Chaetomium thermophilum | genome | http://ct.bork.embl.de/downloads.html | C_thermophilum.annotation.v2.4.tar.gz | 20181107 |
| **1305733** | Rhodotorula sp | genome | ftp://ftp.ensemblgenomes.org/pub/fungi/release-34/fasta/fungi_basidiomycota1_collection/rhodotorula_sp_jg_1b/pep/Rhodotorula_sp_jg_1b.Rhosp1.pep.all.fa.gz | Rhosp1 | 20170120 |
| **28583** | Allomyces macrogynus | genome | ftp://ftp.ensemblgenomes.org/pub/fungi/release-34/fasta/fungi_blastocladiomycota1_collection/allomyces_macrogynus_atcc_38327/pep/Allomyces_macrogynus_atcc_38327.A_macrogynus_V3.pep.all.fa.gz | A_macrogynus_V3 | 20170120 |
| **109876** | Catenaria anguillulae | genome | http://genome.jgi.doe.gov/Catan2/download/Catan2_all_proteins_20160412.aa.fasta.gz | Catan2 | 20170120 |
| **109871** | Batrachochytrium dendrobatidis | genome | ftp://ftp.ensemblgenomes.org/pub/fungi/release-34/fasta/fungi_chytridiomycota1_collection/batrachochytrium_dendrobatidis_jam81/pep/Batrachochytrium_dendrobatidis_jam81.v1.0.pep.all.fa.gz | V1.0 | 20170120 |
| **645134** | Spizellomyces punctatus DAOM | genome | ftp://ftp.ensemblgenomes.org/pub/fungi/release-34/fasta/fungi_chytridiomycota1_collection/spizellomyces_punctatus_daom_br117/pep/Spizellomyces_punctatus_daom_br117.S_punctatus_V1.pep.all.fa.gz | S_punctatis_V1 | 20170120 |
| **423460** | Basidiobolus meristosporus | genome | http://genome.jgi.doe.gov/Basme2finSC/download/Basme2finSC_GeneCatalog_proteins_20150701.aa.fasta.gz | BasmefinSC | 20170120 |

| | | | | | |
|---|---|---|---|---|---|
| **796925** | Condiobolus coronatus | genome | ftp://ftp.ensemblgenomes.org/pub/fungi/release-34/fasta/fungi_entomophthoromycota1_collection/conidiobolus_coronatus_nrrl_28638/pep/Conidiobolus_coronatus_nrrl_28638.Conidiobolus_coronatus_NRRL28638.pep.all.fa.gz | Conidiobolus coronatus NRRL28638 | 20170120 |
| **588596** | Rhizophagus irregularis | genome | ftp://ftp.ensemblgenomes.org/pub/fungi/release-34/fasta/fungi_glomeromycota1_collection/rhizophagus_irregularis_daom_181602/pep/Rhizophagus_irregularis_daom_181602.Gloin1.pep.all.fa.gz | Gloin1 | 20170120 |
| **61392** | Coemansia reversa | genome | genome.jgi.doe.gov/Coere1/download/Coere1_GeneCatalog_proteins_20110909.aa.fasta.gz | Coere1 | 20170120 |
| **304332** | Ramicandelaber brevisporus | genome | http://genome.jgi.doe.gov/Rambr1/download/Rambr1_GeneCatalog_proteins_20140929.aa.fasta.gz | Rambr1 | 20170120 |
| **1485682** | Mitosporidium daphniae | genome | ftp://ftp.ensemblgenomes.org/pub/fungi/release-34/fasta/fungi_microsporidia1_collection/mitosporidium_daphniae/pep/Mitosporidium_daphniae.UGP1.0.pep.all.fa.gz | UGP1.0 | 20170125 |
| **58839** | Encephalitozoon intestinalis | genome | ftp://ftp.ensemblgenomes.org/pub/fungi/release-34/fasta/fungi_microsporidia1_collection/encephalitozoon_intestinalis_atcc_50506/pep/Encephalitozoon_intestinalis_atcc_50506.ASM14646v1.pep.all.fa.gz | ASM14646v1 | 20170125 |
| **78898** | Mortierella verticillata | genome | http://genome.jgi.doe.gov/Morve1/download/Morve1_GeneCatalog_proteins_20161024.aa.fasta.gz | Morve1 | 20170120 |
| **310910** | Mortierella elongata | genome | http://genome.jgi.doe.gov/Morel2/download/Morel2_GeneCatalog_proteins_20151120.aa.fasta.gz | Morel2 | 20170120 |
| **36080** | Mucor circinelloides | genome | http://genome.jgi.doe.gov/Mucci2/download/Mucor_circinelloides_v2_filtered_proteins.fasta.gz | Muccl2 | 20170120 |
| **4837** | Phycomyces blakesleeanus | genome | http://genome.jgi.doe.gov/Phybl2/download/Phycomyces_blakesleeanus_v2_filtered_proteins.fasta.gz | Phybl2 | 20170120 |
| **1577477** | Piromyces sp finn | genome | http://genome.jgi.doe.gov/Pirfi3/download/Pirfi3_GeneCatalog_proteins_20160330.aa.fasta.gz | Pirfi3 | 20170120 |
| **1550276** | Neocallimastix sp | genome | http://genome.jgi.doe.gov/Neosp1/download/Neosp1_GeneCatalog_proteins_20160330.aa.fasta.gz | Neosp1 | 20170120 |
| **5141** | Neurospora crassa | genome | ftp://ftp.ensemblgenomes.org/pub/fungi/release-34/fasta/fungi_ascomycota2_collection/neurospora_crassa_gca_000786625/pep/Neurospora_crassa_gca_000786625.Neucr_trp3_1.pep.all.fa.gz | Neucr_trp3_1 | 20170208 |
| **39416** | Tuber melanosporum | genome | ftp://ftp.ensemblgenomes.org/pub/fungi/release-34/fasta/tuber_melanosporum/pep/Tuber_melanosporum.ASM15164v1.pep.all.fa.gz | ASM15164v1 | 20170208 |
| **242477** | Melampsora laricis | genome | http://genome.jgi.doe.gov/Mellp2_3/download/Mellp2_3_GeneCatalog_proteins_20151130.aa.fasta.gz | mellp2_3 | 20170120 |
| **4932** | Saccharomyces cerevisiae | genome | ftp://ftp.ensemblgenomes.org/pub/fungi/release-34/fasta/saccharomyces_cerevisiae/pep/Saccharomyces_cerevisiae.R64-1-1.pep.all.fa.gz | R64-1-1 | 20170208 |

| | | | | | |
|---|---|---|---|---|---|
| **28985** | Kluyveromyces lactis | genome | ftp://ftp.ensemblgenomes.org/pub/fungi/release-34/fasta/fungi_ascomycota1_collection/kluyveromyces_lactis/pep/Kluyveromyces_lactis.ASM251v1.pep.all.fa.gz | ASM251v1 | 20170208 |
| **4952** | Yarrowia lipolytica | genome | ftp://ftp.ncbi.nlm.nih.gov/genomes/all/GCA/001/761/485/GCA_001761485.1_ASM176148v1/GCA_001761485.1_ASM176148v1_protein.faa.gz | ASM176148v1 | 20170208 |
| **5606** | Saitoella complicata | genome | ftp://ftp.ensemblgenomes.org/pub/fungi/release-34/fasta/fungi_ascomycota2_collection/saitoella_complicata_nrrl_y_17804/pep/Saitoella_complicata_nrrl_y_17804.Scomplicata_3.0.pep.all.fa.gz | Scomplicata_3.0 | 20170120 |
| **4896** | Schizosaccharomyces pombe | genome | ftp://ftp.ensemblgenomes.org/pub/fungi/release-34/fasta/schizosaccharomyces_pombe/pep/Schizosaccharomyces_pombe.ASM294v2.pep.all.fa.gz | ASM294v2 | 20170120 |
| **5270** | Ustilago maydis | genome | ftp://ftp.ensemblgenomes.org/pub/fungi/release-34/fasta/ustilago_maydis/pep/Ustilago_maydis.Umaydis521_2.0.pep.all.fa.gz | Umaydis521_2.0 | 20170120 |
| **1851185** | Syncephalis plumigaleata | genome | http://genome.jgi.doe.gov/Synplu1/download/Synplu1_primary_alleles_proteins_20160908.aa.fasta.gz | synplu1 | 20170120 |
| **281847** | Rozella allomycis | genome | ftp://ftp.ensemblgenomes.org/pub/fungi/release-34/fasta/fungi_rozellomycota1_collection/rozella_allomycis_csf55/pep/Rozella_allomycis_csf55.Rozella_k41_t100.pep.all.fa.gz | Rozella_k41_t100 | 20170120 |
| **749232** | Abeoforma whisleri | genome | https://figshare.com/articles/Genome_-_Abeoforma_whisleri_/5426458 | | 20181111 |
| **1932427** | Chromosphaera perkinsii | genome | https://figshare.com/articles/Genome_-_Chromosphaera_perkinsii/5426494 | | 20181111 |
| **39843** | Ichthyophonus hoferi | genome | https://figshare.com/articles/Genome_-_Ichthyophonus_hoferi/5426488 | | 20181111 |
| **749231** | Pirum gemmata | genome | https://figshare.com/articles/Genome_-_Pirum_gemmata/5426506 | | 20181111 |
| **72019** | Sphaeroforma arctica | genome | https://figshare.com/articles/dataset/Sphaeroforma_arctica_transcriptome/8299529 , , 20220920 | Sarc4 | 20220920 |
| **470921** | Creolimax fragrantissima | genome | https://figshare.com/articles/Creolimax_fragrantissima_genome_data/1403592 | Creollimax_fragrantissima | 20170120 |
| **1553916** | Nuclearia sp | transcriptome | https://figshare.com/articles/Nuclearia_sp_ATCC_50694_-_Transcriptome/3898485 | Nuclearia_a_unigene | 20170120 |
| **691883** | Fonticula alba | genome | ftp://ftp.ncbi.nlm.nih.gov/genomes/all/GCA/000/388/065/GCA_000388065.2_Font_alba_ATCC_38817_V2/GCA_000388065.2_Font_alba_ATCC_38817_V2_protein.faa.gz | Font_alba_ATCC_38817_V2 | 20170120 |
| **2717461** | Colponemidia sp Colp-10 | transcriptome | https://sra-download.ncbi.nlm.nih.gov/traces/wgs01/wgs_aux/GI/LJ/GILJ01/GILJ01.1.fsa_nt.gz | GILJ01000000 | 20210527 |
| **2717462** | Colponemidia sp Colp-15 | transcriptome | https://sra-download.ncbi.nlm.nih.gov/traces/wgs01/wgs_aux/GI/LK/GILK01/GILK01.1.fsa_nt.gz | GILK01000000 | 20210527 |

| | | | | | |
|---|---|---|---|---|---|
| **5833** | Plasmodium falciparum | genome | ftp://ftp.ensemblgenomes.org/pub/release-34/protists/fasta/plasmodium_falciparum/pep/ | ASM276v1 | 20161212 |
| **5866** | Babesia bigemina | genome | ftp://ftp.ensemblgenomes.org/pub/release-34/protists/fasta/protists_alveolata1_collection/babesia_bigemina/pep/ | Bbig001 | 20161212 |
| **5874** | Theileria annulata | genome | ftp://ftp.ensemblgenomes.org/pub/release-34/protists/fasta/protists_alveolata1_collection/theileria_annulata/pep/ | ASM322v1 | 20161212 |
| **110365** | Gregarina niphandrodes | genome | ftp://ftp.ensemblgenomes.org/pub/release-34/protists/fasta/protists_alveolata1_collection/gregarina_niphandrodes/pep/ | GNI3 | 20161212 |
| **5808** | Cryptosporidium muris | genome | ftp://ftp.ensemblgenomes.org/pub/release-34/protists/fasta/protists_alveolata1_collection/cryptosporidium_muris_rn66/pep/ | JCVI-cmg-v1.0 | 20161212 |
| **5807** | Cryptosporidium parvum | genome | ftp://ftp.ensemblgenomes.org/pub/release-34/protists/fasta/protists_alveolata1_collection/cryptosporidium_parvum_iowa_ii/pep/ | ASM16534v1 | 20161212 |
| **5811** | Toxoplasma gondii | genome | ftp://ftp.ensemblgenomes.org/pub/release-34/protists/fasta/toxoplasma_gondii/pep/ | ToxoDB-7.1 | 20161212 |
| **5801** | Eimeria acervulina | genome | ftp://ftp.ensemblgenomes.org/pub/release-34/protists/fasta/protists_alveolata1_collection/eimeria_acervulina/pep/ | EAH001 | 20161212 |
| **505693** | Chromera velia | genome | http://cryptodb.org/common/downloads/Current_Release/CveliaCCMP2878/fasta/data/ | ? | 20170208 |
| **1169539** | Vitrella brassicaformis | genome | ftp://ftp.ensemblgenomes.org/pub/release-34/protists/fasta/protists_alveolata1_collection/vitrella_brassicaformis_ccmp3155/pep/ | Vbrassicaformis | 20161212 |
| **5888** | Paramecium tetraurelia | genome | ftp://ftp.ensemblgenomes.org/pub/release-34/protists/fasta/paramecium_tetraurelia/pep/ | GCA_000165425.1 | 20161212 |
| **5911** | Tetrahymena thermophila | genome | ftp://ftp.ensemblgenomes.org/pub/release-34/protists/fasta/tetrahymena_thermophila/pep/ | JCVI-TTA1-2.2 | 20161212 |
| **5932** | Ichthyophthirius multifiliis | genome | ftp://ftp.ensemblgenomes.org/pub/release-34/protists/fasta/protists_alveolata1_collection/ichthyophthirius_multifiliis/pep/ | JCVI-IMG1-V.1 | 20161212 |
| **266149** | Pseudocohnilembus persalinus | genome | ftp://ftp.ensemblgenomes.org/pub/release-34/protists/fasta/protists_alveolata1_collection/pseudocohnilembus_persalinus/pep/ | ASM144751v1 | 20161212 |
| **5949** | Stylonychia lemnae | genome | ftp://ftp.ensemblgenomes.org/pub/release-34/protists/fasta/protists_alveolata1_collection/stylonychia_lemnae/pep/ | Stylonychia_lemnae_asm_v1.0 | 20161212 |
| **1172189** | Oxytricha trifallax | genome | ftp://ftp.ensemblgenomes.org/pub/release-34/protists/fasta/protists_alveolata1_collection/oxytricha_trifallax_gca_000295675/ | GCA_00295675 | 20161212 |
| **5963** | Stentor coeruleus | genome | https://www.ncbi.nlm.nih.gov/assembly/GCA_001970955.1/ | ASM197095v1 \| S_coeruleus_Nov216 | 20170210 |

| | | | | | |
|---|---|---|---|---|---|
| **31276** | Perkinsus marinus | genome | ftp://ftp.ensemblgenomes.org/pub/release-34/protists/fasta/protists_alveolata1_collection/perkinsus_marinus_atcc_50983/pep/ | JCVI_PMG_1.0 | 20161212 |
| **2951** | Symbiodinium microadriaticum | genome | ftp://ftp.ncbi.nlm.nih.gov/genomes/all/GCA/001/939/145/GCA_001939145.1_ASM193914v1/GCA_001939145.1_ASM193914v1_protein.faa.gz | ASM193914v1 | 20170207 |
| **1202447** | Symbiodinium minutum | genome | http://marinegenomics.oist.jp/symb/viewer/download?project_id=21 (Assembly V1.0, symbB.v1.2.augustus.prot.fa.gz) | Symbiodinium minutum ver. symb_aug_v1.120123 | 20161212 |
| **37360** | Plasmodiophora brassicae | genome | ftp://ftp.ensemblgenomes.org/pub/release-34/protists/fasta/protists_rhizaria1_collection/plasmodiophora_brassicae/pep/ | pbe3.h15 | 20161212 |
| **46433** | Reticulomyxa filosa | genome | ftp://ftp.ensemblgenomes.org/pub/release-34/protists/fasta/protists_rhizaria1_collection/reticulomyxa_filosa/pep/ | Reti_assembly1.0 | 20161212 |
| **227086** | Bigelowiella natans | transcriptome | https://figshare.com/articles/Marine_Microbial_Eukaryotic_Transcriptome_Sequencing_Project_re-assemblies/3840153/3 | version 3 | 20181113 |
| **552665** | Bigelowiella longifila | transcriptome | https://figshare.com/articles/Marine_Microbial_Eukaryotic_Transcriptome_Sequencing_Project_re-assemblies/3840153/3 | version 3 | 20181113 |
| **29199** | Chlorarachnion reptans | transcriptome | https://figshare.com/articles/Marine_Microbial_Eukaryotic_Transcriptome_Sequencing_Project_re-assemblies/3840153/3 | version 3 | 20181113 |
| **2850** | Phaeodactylum tricornutum | genome | ftp://ftp.ensemblgenomes.org/pub/release-34/protists/fasta/phaeodactylum_tricornutum/pep/ | ASM15095v2 | 20161212 |
| **35128** | Thalassiosira pseudonana | genome | ftp://ftp.ensemblgenomes.org/pub/release-34/protists/fasta/thalassiosira_pseudonana/pep/ | ASM14940v2 | 20161212 |
| **186039** | Fragilariopsis cylindrus | genome | http://genome.jgi.doe.gov/pages/dynamicOrganismDownload.jsf?organism=Fracy1 (Fracy1_GeneModels_FilteredModels1_aa.fasta.gz) | v1.0 | 20170105 |
| **72520** | Nannochloropsis gaditana | genome | http://www.nannochloropsis.org/page/ftp | CCMP526 | 20170327 |
| **87111** | Aplanochytrium kerguelense | genome | http://genome.jgi.doe.gov/pages/dynamicOrganismDownload.jsf?organism=Aplke1 (Aplke1_GeneCatalog_proteins_20121220.aa.fasta.gz) | V1.0 | 20161212 |
| **4773** | Schizochytrium aggregatum | genome | http://genome.jgi.doe.gov/pages/dynamicOrganismDownload.jsf?organism=Schag1 (Schag1_GeneCatalog_proteins_20121220.aa.fasta.gz) | V1.0 | 20161212 |
| **87102** | Aurantiochytrium limacinum | genome | http://genome.jgi.doe.gov/pages/dynamicOrganismDownload.jsf?organism=Aurli1 (Aurli1_GeneCatalog_proteins_20120618.aa.fasta.gz) | V1.0 | 20161212 |
| **4787** | Phytophthora infestans | genome | ftp://ftp.ensemblgenomes.org/pub/release-34/protists/fasta/phytophthora_infestans/pep/ | ASM14294v1 | 20161212 |
| **123356** | Hyaloperonospora parasitica | genome | ftp://ftp.ensemblgenomes.org/pub/release-34/protists/fasta/hyaloperonospora_arabidopsidis/pep/ | HyaAraEmoy2_2.0 | 20161212 |

| | | | | | |
|---|---|---|---|---|---|
| **4781** | Plasmopara halstedii | genome | ftp://ftp.ensemblgenomes.org/pub/release-34/protists/fasta/protists_stramenopiles1_collection/plasmopara_halstedii/pep/ | | 20161212 |
| **2052682** | Pythium ultimum | genome | ftp://ftp.ensemblgenomes.org/pub/release-34/protists/fasta/pythium_ultimum/pep/ | | 20161212 |
| **653948** | Albugo laibachii | genome | ftp://ftp.ensemblgenomes.org/pub/release-34/protists/fasta/albugo_laibachii/pep/ | ENA 1 | 20161212 |
| **112090** | Aphanomyces astaci | genome | ftp://ftp.ensemblgenomes.org/pub/release-34/protists/fasta/protists_stramenopiles1_collection/aphanomyces_astaci/pep/ | Apha_asta_APO3_V1 | 20161212 |
| **101203** | Saprolegnia parasitica | genome | ftp://ftp.ensemblgenomes.org/pub/release-34/protists/fasta/protists_stramenopiles1_collection/saprolegnia_parasitica_cbs_223_65/pep/ | ASM15154v2 | 20161212 |
| **12968** | Blastocystis hominis | genome | ftp://ftp.ensemblgenomes.org/pub/release-34/protists/fasta/protists_stramenopiles1_collection/blastocystis_hominis/pep/ | ASM15166v1 | 20161212 |
| **44056** | Aureococcus anophagefferens | genome | ftp://ftp.ensemblgenomes.org/pub/release-34/protists/fasta/protists_stramenopiles1_collection/aureococcus_anophagefferens/pep/ | v 1.0 | 20161212 |
| **2880** | Ectocarpus siliculosus | genome | https://bioinformatics.psb.ugent.be/gdb/ectocarpus/ (EctsiV2_prot_LATEST.tfa.gz) | EctsiV2 | 20161212 |
| **309737** | Cladosiphon okamuranus | genome | http://marinegenomics.oist.jp/algae/viewer/download?project_id=53 (downloaded file: 160208_2k_oki_prot.fa.gz) | Assembly v1.0 | 20161212 |
| **529818** | Thecamonas trahens | genome | ftp://ftp.ensemblgenomes.org/pub/protists/release-34/fasta/protists_apusozoa1_collection/thecamonas_trahens_atcc_50062/pep/Thecamonas_trahens_atcc_50062.TheTra_May2010.pep.all.fa.gz | TheTra_May2010 | 20170120 |
| **221724** | Seculamonas sp ecuadoriensis | EST | https://www.ncbi.nlm.nih.gov/nuccore?LinkName=biosample_nuccore&from_uid=150550 | LIBEST_019958 | 20210527 |
| **392300** | Histiona aroides | EST | https://www.ncbi.nlm.nih.gov/nuccore?LinkName=biosample_nuccore&from_uid=150562 | LIBEST_019970 | 20210527 |
| **221721** | Jakoba bahamiensis | EST | https://www.ncbi.nlm.nih.gov/nuccore?LinkName=biosample_nuccore&from_uid=150510 | LIBEST_019913 | 20210527 |
| **143017** | Jakoba libera | EST | https://www.ncbi.nlm.nih.gov/nuccore?LinkName=biosample_nuccore&from_uid=150512 | LIBEST_019915 | 20210527 |
| **48483** | Reclinomonas americana | EST | https://www.ncbi.nlm.nih.gov/nuccore?LinkName=biosample_nuccore&from_uid=150540,https://www.ncbi.nlm.nih.gov/nuccore?LinkName=biosample_nuccore&from_uid=150541,https://www.ncbi.nlm.nih.gov/nuccore?LinkName=biosample_nuccore&from_uid=150542,https://www.ncbi.nlm.nih.gov/nuccore?LinkName=biosample_nuccore&from_uid=150543,https://www.ncbi.nlm.nih.gov/nuccore?LinkName=biosample_nuccore&from_uid=150544,https://www.ncbi.nlm.nih.gov/nuccore?LinkName=biosample_nuccore&from_uid=150545,https://www.ncbi.nlm.nih.gov/nuccore?LinkName=biosample_nuccore&from_uid=150546 | LIBEST_019948,LIBEST_019949,LIBEST_019950,LIBEST_019951,LIBEST_019952,LIBEST_019953, LIBEST_019954 | 20210527 |

| | | | | | |
|---|---|---|---|---|---|
| **505711** | Andalucia godoyi | genome | https://megasun.bch.umontreal.ca/Andalucia_godoyi/Andalucia_godoyi_proteome.faa | Andalucia_godoyi_proteome | 20210527 |

**Supplementary text**

*Dam1-C subunits are not syntenic*

While collecting Dam1-C homologs also their genomic localization was analyzed because (especially recent) transfer in their evolution could leave traces like co-localization in the genome. For 18 out of the 32 species, the genes encoding subunits of the Dam1-C were located on different scaffolds/chromosomes, while in 13 of the 32 species, 2 or 3 of the subunits are on the same scaffold/chromosome, but separated by on average ±560kb nucleotides. In the case that HGT (Horizontal Gene Transfer) explains the alternating pattern between Ska-C and Dam1-C, would be due to HGT, one would expect that this happened in a single HGT event comprising all Dam1-C subunits, enabled by and resulting in their close genomic eukaryotic tree. The current-day genomic localization of Dam1-c subunits is not consistent with enabling nor receiving HGT.
